# Combined behavioral and electrophysiological evidence for a direct cortical effect of prefrontal tDCS on disorders of consciousness

**DOI:** 10.1101/612309

**Authors:** Bertrand Hermann, Federico Raimondo, Lukas Hirsch, Yu Huang, Mélanie Denis-Valente, Pauline Pérez, Denis Engemann, Frédéric Faugeras, Nicolas Weiss, Sophie Demeret, Benjamin Rohaut, Lucas C. Parra, Jacobo D. Sitt, Lionel Naccache

## Abstract

Severe brain injuries can lead to long-lasting disorders of consciousness (DoC) such as vegetative state/unresponsive wakefulness syndrome (VS/UWS) or minimally conscious state (MCS). While behavioral assessment remains the gold standard to determine conscious state, EEG has proven to be a promising complementary tool to monitor the effect of new therapeutics. Encouraging results have been obtained with invasive electrical stimulation of the brain, and recent studies identified transcranial direct current stimulation (tDCS) as an effective approach in randomized controlled trials. This non-invasive and inexpensive tool may turn out to be the preferred treatment option. However, its mechanisms of action and physiological effects on brain activity remain unclear and debated. Here, we stimulated 60 DoC patients with the anode placed over left-dorsolateral prefrontal cortex in a prospective open-label study. Clinical behavioral assessment improved in twelve patients (20%) and none deteriorated. This behavioral response after tDCS coincided with an enhancement of putative EEG markers of consciousness: in comparison with non-responders, responders showed increases of power and long-range cortico-cortical functional connectivity in the theta-alpha band, and a larger and more sustained P300 suggesting improved conscious access to auditory novelty. The EEG changes correlated with electric fields strengths in prefrontal cortices, and no correlation was found on the scalp. Taken together, this prospective intervention in a large cohort of DoC patients strengthens the validity of the proposed EEG signatures of consciousness, and is suggestive of a direct causal effect of tDCS on consciousness.

## MAIN TEXT

### Introduction

After an acute phase of coma, severe acute brain injuries can lead to lasting disorders of consciousness (DoC). The Coma Recovery Scale-Revised (CRS-R) (1, 2) is the most widely accepted tool to distinguish vegetative state/unresponsive wakefulness syndrome (VS/UWS) from minimally conscious state (MCS) patients. While VS/UWS only show non-purposeful reflexive behaviors, MCS show reproducible yet inconsistent cognitive and intentional cortically-mediated behaviors (3). Finally, exit-MCS patients exhibit functional communication or use of objects. However, this behavioral assessment has limitations and 15-20% of behaviorally-diagnosed VS/UWS patients show patterns of brain activity suggestive of higher states of consciousness (4–8). Among the different brain-imaging techniques used, EEG has proved to be a reliable, non-invasive bedside tool to probe signatures of both conscious state and conscious access to external stimuli in DoC patients (9–15). Specifically, increases of spectral power, complexity and functional connectivity in the theta-alpha bands correlate with state of consciousness, but the specificity and causal value of these putative signatures of consciousness remain to be demonstrated. The combination of such behavioral and EEG measures seems optimal to assess in detail possible improvements of consciousness during treatment with new therapeutic techniques.

Encouraging results have been obtained on DoC patients with invasive electrical stimulation of the brain (16–19) and recently, transcranial delivery of a low intensity electrical current over the scalp with tDCS (20–22). However, the efficacy of tDCS is debated (23–25), and its mechanisms have not been established yet.

In this prospective case-control open-label study, we evaluated the impact of prefrontal tDCS (Figure 1A) on both the behavior and quantified electrophysiological measures of conscious state and conscious access. We then investigated the possible mechanism of action of tDCS by correlating the electrophysiological response to the applied electric fields estimated from individual patients’ head and brain MRI anatomy.

**Figure 1.**
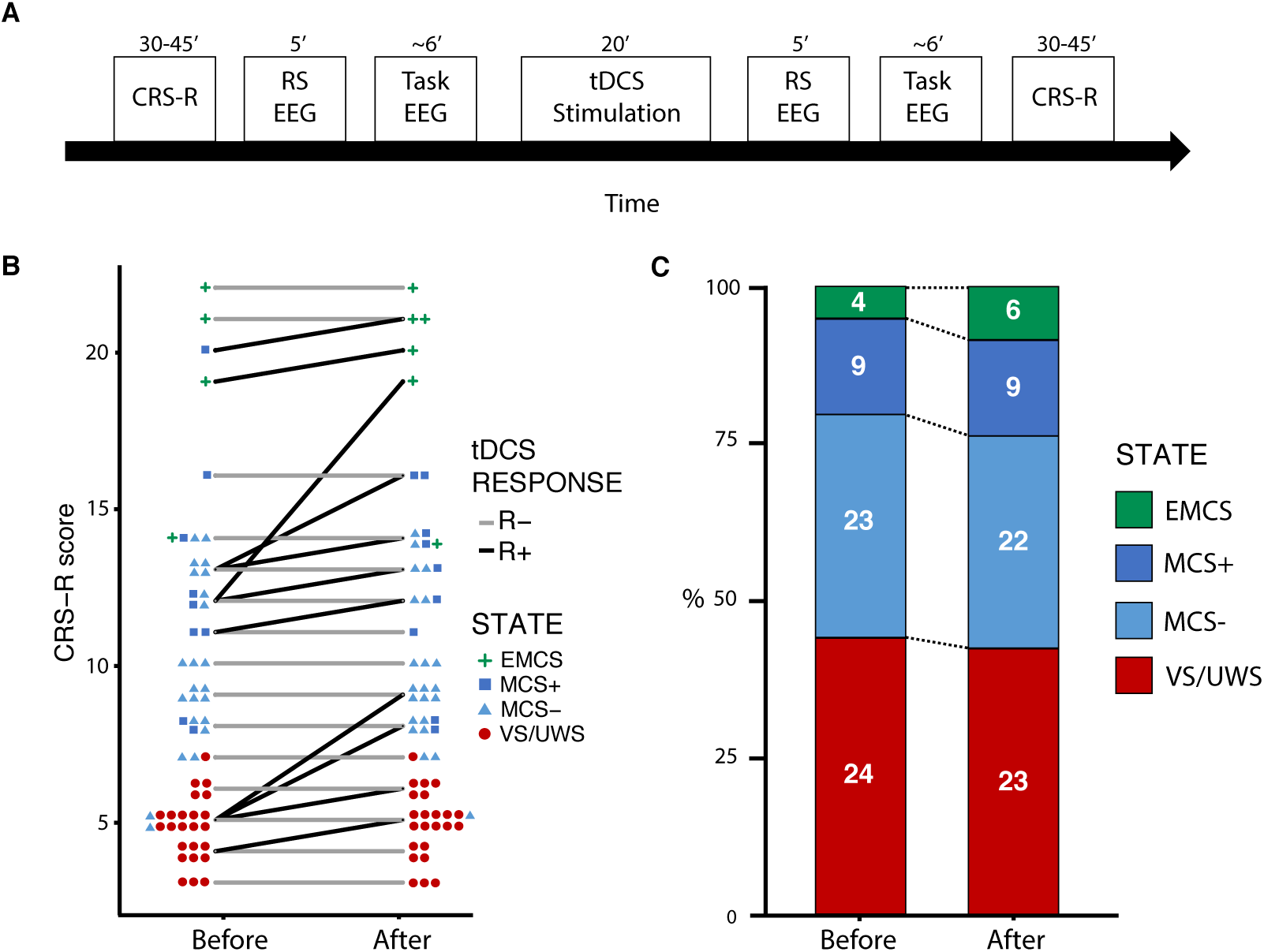
Behavioral response to tDCS. (A) Study protocol timeline showing the behavioral (Coma Recovery Scale-revised (CRS-R) and electrophysiological (resting-state (RS) and auditory oddball paradigm (task-EEG)) measures of the effects consecutive to a single transcranial direct current stimulation (tDCS) session. (B) Individual patients’ CRS-R scores before and after tDCS are represented for responders (R+, in black) and non-responders (R-, in gray), together with the number of patients and their state (symbols). (C) The proportion of each state of consciousness, before and after tDCS showed an increase in the higher states of consciousness (exit minimally conscious state (EMCS) and minimally conscious state ‘plus’ (MCS+)), at the expense of lower states of consciousness (vegetative state/unresponsive wakefulness syndrome (VS/UWS) and MCS ‘minus’ (MCS-)).

### Results

#### Behavioral response after one tDCS session

Between October 2015 and September 2018, among 69 eligible DoC patients, 66 patients were treated prospectively with a single 20 minutes tDCS session with the anode placed over the left dorsolateral prefrontal cortex and the cathode over the right supraorbital cortex same parameters as previously reported in DoC patients (21). The effects of this tDCS session were evaluated by a combined behavioral and electrophysiological assessment using CRS-R and high-density EEG recordings at rest and during an auditory oddball paradigm immediately before and after the stimulation (Figure 1). Among these, 60 patients could be included in the study (one was comatose, one had a seizure during EEG and four had insufficient EEG data for subsequent analyses, *SI section B1*, Figure S1). This cohort was composed of 24 VS/UWS, 32 MCS and 4 exit-MCS patients (*SI section B1*, Table S1). Response to tDCS (R+), defined as an increase of the CRS-R score after stimulation as compared to before, was observed in 12 patients (20%): 4 VS/UWS (16.7%), 7 MCS (21.9%) and 1 exit-MCS patients (25%), without differences across groups (*p*>0.8, Fisher’s exact test). This resulted in a change in consciousness state in 3 patients, one VS/UWS shifting towards MCS and two MCS shifting towards exit-MCS (Figure 1B and 1C). This proportion of R+ is close to the 27% reported in a double-blind randomized trial with the same stimulation parameters (21). Conversely, non-responders (R-) were defined by CRS-R scores that were either stable or decreasing after tDCS. Interestingly, none of the patients showed a decrease of CRS-R score. Overall, CRS-R score changes after stimulation were significantly larger than zero (*p=*0.002, Wilcoxon signed-rank test, effect size *r=*0.28 [0.21-0.36]). No significant differences were found between R+ and R-populations in their demographic characteristics (age, sex, etiology and time since injury), and, importantly, neither in their vigilance before tDCS, nor data preprocessing (*SI section B1*, Table S2).

#### Spectral power and connectivity in the theta-alpha band increase in responders to tDCS

We first analyzed the interaction between resting-state brain activity and the behavioral response after tDCS stimulation using 5-minutes of EEG acquired before and after stimulation. To that end, we computed putative signatures of conscious state on both recordings using an automated procedure reported previously ((26), see Methods). We then tested whether these EEG signatures were modified by tDCS differently in R+ than in R-, using the following contrast: [post – pre] in R+ > [post – pre] in R-.

*Power Spectral Density.* As compared to R-, R+ showed a significant increase after the stimulation in normalized theta power with a topography maximal over the parietal cortices (*p*=0.03, effect size *g*=0.92 [0.56-2.03]; Figure 2). Similarly, we observed an increase of both raw and normalized alpha power (*p*=0.01, *g*=0.80 [0.44-0.80] and *p*=0.04, *g*=1.10 [0.79-1.91] respectively). Other markers of spectral power did not differ between R- and R+ (*SI section B2*, Figure S2).

**Figure 2.**
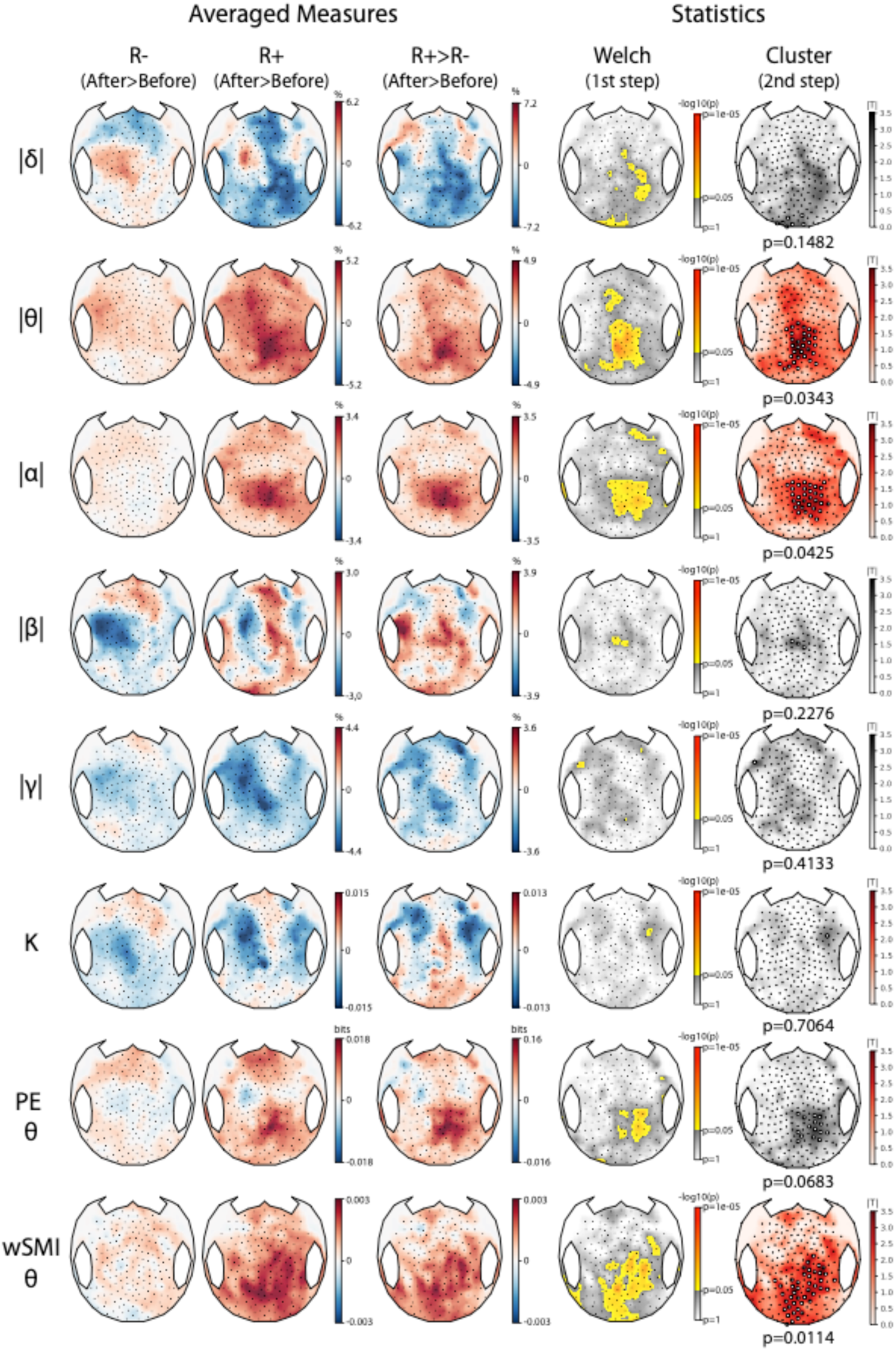
Resting state EEG markers increases after tDCS in responders. Topographic representations of the tDCS-induced changes in normalized spectral power (delta |δ|, theta |θ|, alpha |α|, beta |β| and gamma |γ|), Kolmogorov complexity (K), permutation entropy in the theta-alpha band (PE θ) and weighted symbolic mutual information in the theta-alpha band (wSMI θ) over the 224 scalp electrodes according to the behavioral response to tDCS. After minus before differences are presented for both non-responders (R-) and responders (R+) (left columns), followed by the contrast between the two (middle columns) and the corresponding statistical comparison using a two-steps spatial cluster-based permutation approach (right columns). Significant centro-parietal clusters were found for |θ| and |α| power (*p*=0.0343 and *p*=0.0425) and for wSMI θ (*p*=0.0114). Absolute t-values are plotted with a red color scale when a significant cluster was found and in grey otherwise. Electrodes forming the cluster are highlighted by white circles.

*Complexity.* EEG complexity, assessed by the permutation entropy in the theta-alpha band, showed a trend of an increase in R+ as compared to R-in the same parietal region (*p*=0.07, *g*=0.70 [−0.11-1.05]). Kolmogorov and spectral entropy did not differ.

*Functional connectivity.* Response to tDCS was also characterized by an increase of functional connectivity in the theta-alpha band (4-10 Hz), assessed by the weighted symbolic mutual information. When comparing topographies of averaged values, we found a parieto-occipital cluster with increased values in R+ as compared to R-patients (*p*=0.01, *g*=0.82 [0.11-1.62]) (Figure 2, last row). This was confirmed by the analysis conducted across all pairs of electrodes: 4 significant clusters of electrodes located over parietal and occipital cortices showed larger values of functional connectivity for R+ than R-patients (respective p-values were 0.01, 0.02, 0.03 and 0.04 with a global *g*=1.27 [0.52-1.54]; Figure 3A). These differences between R+ and R-were explained by an increase after the stimulation in R+ patients (*p*=0.02, *g*=0.81 [0.54-1.21]), while no significant change was found in R-patients (Figure 3B). We found no differences between R+ and R-in functional connectivity.

**Figure 3.**
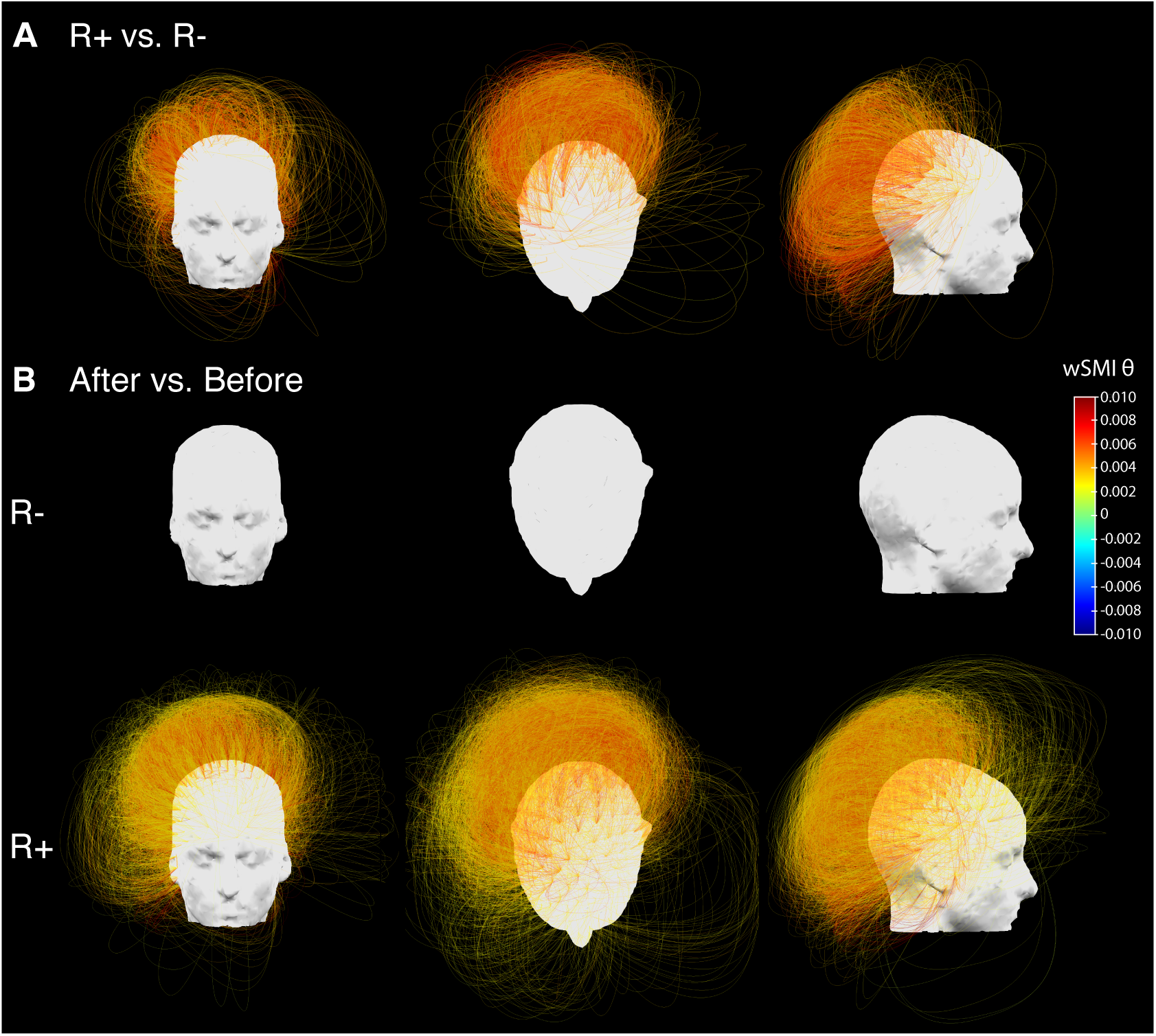
Functional connectivity in the theta-alpha band. Three-dimensional representation of functional connectivity in the theta band assessed by the weighted symbolic mutual information (wSMI θ) showing a significant increase in the centro-posterior regions in responders (R+) compared to non-responders (R-). Four significant clusters involving respectively 902 (*p*=0.01), 438 (*p*=0.02) 363 (*p*=0.03) and 245 (*p*=0.04) pairs of electrodes were identified. For visual clarity, they are plotted together (total of 1948 pairs) (A). Restricted contrast revealed a significant increase in the wSMI θ over centro-parietal regions (one single cluster of 5918 pairs of electrodes, *p*=0.02, see B bottom row) after tDCS in responders, whereas no change could be detected in non-responders (B, upper row).

Note that tDCS effects did not appear systematically different for any of these markers when comparing VS/UWS patients to MCS and exit-MCS patients (*SI section B3*, Figure S3).

#### A neural signature of conscious access improves in responders to tDCS

In addition to resting state, we assessed the impact of tDCS on the ability of patients to detect auditory regularities during an oddball paradigm (modified from (27), a task known to require conscious access to auditory novelty. Patients were instructed to actively count the occurrence of auditory oddballs (series of 4 identical tones followed by a 5^th^ distinct tone; 20% of trials) delivered randomly among series of 5 identical tones (standard trials; 80% of trials). We computed the event-related potentials (ERP) to deviant tone minus standard tone, before and after tDCS, and compared R+ with R-using the same interaction contrast as for the resting state. Five EEG recordings were discarded after automatic assessment of data quality. The analysis performed on the 55 remaining datasets (11 R+ and 44 R-) revealed a significant positive and left-lateralized anterior cluster spanning from 28ms to 376ms after the onset of the 5^th^ sound (*p*=0.008, *g*=1.40 [0.84-1.93], Figure 4A). Pre/post comparisons localized the origin of this effect to two significant clusters in R+ patient (a first posterior cluster from 52-312 ms, *p*=0.03, *g*=-0.96 [−1.52-−0.38] and a second left-lateralized anterior cluster from 68-392 ms, *p*=0.02; *g*=1.26 [0.78-1.84]) while no difference was observed in R-patients (Figure 4B and *SI section B5*, Figure S5A). Note also that although this ERP response started early after the 5^th^ sound it was sustained in time and peaked around 200 ms as a typical P3a component (28, 29).

We supplemented this univariate analysis with a multivariate temporal generalization decoding method (30) in order to better characterize the dynamics of ERP independently from their spatial distribution. This is particularly helpful in studies involving brain-injured patients in which the identification of classical ERP components can fail due to a substantial between-subjects spatial variability. A significant difference was observed as an increase of decoding performances in R+ versus R-patients (two significant clusters with *p*=0.002 and *p*=0.04 respectively, *r*=0.59 [0.07-0.79]; Figure 4C). While R-patients did not show significant clusters when comparing before and after tDCS recordings, R+ patients did show a significant increase of trial class decoding, corresponding to two late and sustained clusters (∼300-600 ms after fifth sound onset; both *p<0.04*; *r*=0.78 [0.61-0.85]; Figure 4D and *SI section B5*, Figure S5B). Notably, this increase of decoding performance after tDCS assumes a square shape from around 300 to 600 ms on the generalization matrix, suggesting a metastable underlying brain activity in this time window (31, 32), evocative of the late P3b signature of conscious access to the violation of auditory novelty (33).

**Figure 4.**
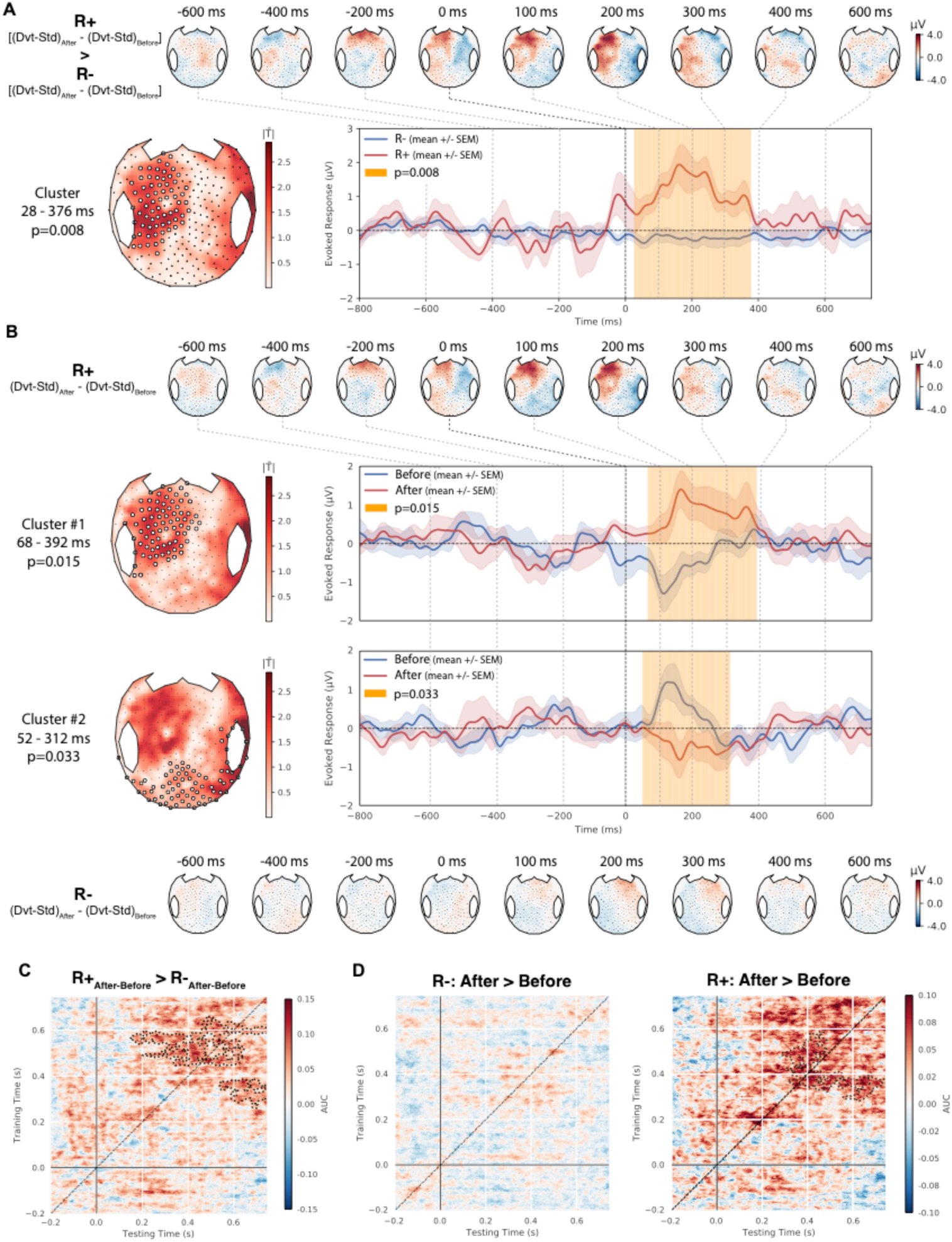
Neural signatures of conscious access to auditory stimuli increases after tDCS in responders. (A). Dynamics of event-related potentials elicited by tDCS in an auditory oddball paradigm (After > Before difference of the Deviant (Dvt) > Standard (Std) contrast), in non-responders (R-, top) and responders (R+, bottom) respectively. (B) A significant spatio-temporal cluster was observed over left fronto-temporal electrodes (white circles) between 28 ms to 376 ms (*p*=0.008; left panel), and the time-course of its voltage amplitude is shown. in R-(blue) and R+ (red). (C) Temporal generalization decoding analysis revealed A significant increase of the decoding performances (after minus before mean AUC) in response to tDCS in R+ as compared to R-, with two significant clusters approximately maximal around 300 ms and 600 ms respectively (*p*=0.002 and *p*=0.04). (D) Restricted comparisons showed that while a significant increase in decoding could be observed in R+ (significant clusters around 300 and 600 ms, *p*=0.03 and *p*=0.04), no such effect could be found in R-. The metastable square pattern in this late time-window is suggestive of an increased P3b component induced by tDCS in R+.

Taken together our results show that clinical behavioral assessment of response to tDCS was characterized by an all-or-none difference: while the R+ group showed a significant effect including a late P3 signature of conscious access to violations of auditory regularities, no such responses could be detected in the R-group, neither with a univariate nor with a multivariate measure.

#### Multivariate prediction of conscious state increases in responders to tDCS

Beyond univariate measures, we also assessed if a behavioral response to tDCS was associated with an improvement in a multivariate EEG-based prediction of conscious state. To that end, we used the support vector machine classifier previously reported to distinguish VS/UWS from MCS using 68 resting-state EEG features (averages and fluctuations over time and space of the univariate markers above). This algorithm was trained on a previously published database of 142 EEG recordings obtained from 98 patients (75 VS/UWS and 67 MCS) (13). For each patient we computed the prediction of being classified MCS, before and after tDCS, and used a nonparametric analysis of repeated measure factorial design with MCS prediction as the dependent variable, the behavioral response as between-subject factor (R+ vs. R-), and stimulation as within-subject factor (pre-vs. post-tDCS). While no main effect of either stimulation or behavioral response was observed, a significant stimulation by behavioral response interaction was present (F(1, 58)=4.2, p=0.045). Post-hoc testing with Wilcoxon signed-rank test showed that while a significant increase of MCS prediction after tDCS was present in R+ patients (median difference of 5.0 % [0.9; 13.7], p=0.01, r=0.51 [0.19-0.63), no effect was found in R-patients (median difference of 2.5 % [−5.3; 7.6], p=0.32, r=0.10 [0.0-0.29]).

#### Electrophysiological response correlates with electric fields intensity in prefrontal cortices

So far, we showed that behavioral improvement of consciousness after tDCS was paralleled by an enrichment of objective EEG measures of both conscious state and conscious access. In order to assess whether this enrichment of EEG activity was specifically mediated by tDCS, - and if so, by which mechanisms -, we then investigated the relation between EEG changes and tDCS-induced electric fields.

Several mechanisms of action of tDCS have been proposed, including modulation of neuronal excitability (34) and effects on synaptic plasticity (35). These two mechanisms, which have been conclusively demonstrated in in-vitro and in-vivo animal studies are driven by the polarization of the neuronal membrane that is brought about by the applied electric field. This polarization increases linearly with field strength (34) and it is therefore reasonable to assume that effects on neural function scale with electric field magnitude. There are some concerns that conventional protocols are sufficient in terms of field intensity to affect neuronal circuits (36), and more recently there also have been suggestions that the effects of transcranial stimulation may result from the activation of peripheral nerves and in particular scalp sensations which have been well-documented (37). These peripheral effects also scale with electric field magnitude by virtue of the same membrane polarization of peripheral nerves. To determine the possible mechanism of action of the stimulation in our population, and disentangle cortical vs. peripheral effects, we analyzed the relation between electric fields in brain and scalp on the one hand and EEG effects on the other.

To that end, we estimated tDCS-induced electric fields in the entire head of patients, based on available T1-weighted MRI (n=47). We then correlated electric fields across patients with the pre/post difference in EEG effects. For EEG we used the multivariate predictor of conscious state described in the previous section. Correlation analysis was limited to areas with mean electric fields likely to have a physiological effect, i.e. >0.5 V/m (38, 39). Areas with positive correlations included both left-dorsolateral and supraorbital prefrontal cortices close to stimulating electrodes (*r*=0.433, *p*=0.007, permutation test), while negative correlations were exclusively confined to the adjacent scalp (*r*=-0.432, *p*=0.036, permutation test, Figure 5). It is hard to reconcile a peripheral effect of skin sensation with negative correlations; whereby stronger EEG effects coincide with weaker sensations. Instead, the results suggest that stronger fields in frontal cortical areas improved EEG markers of consciousness.

**Figure 5.**
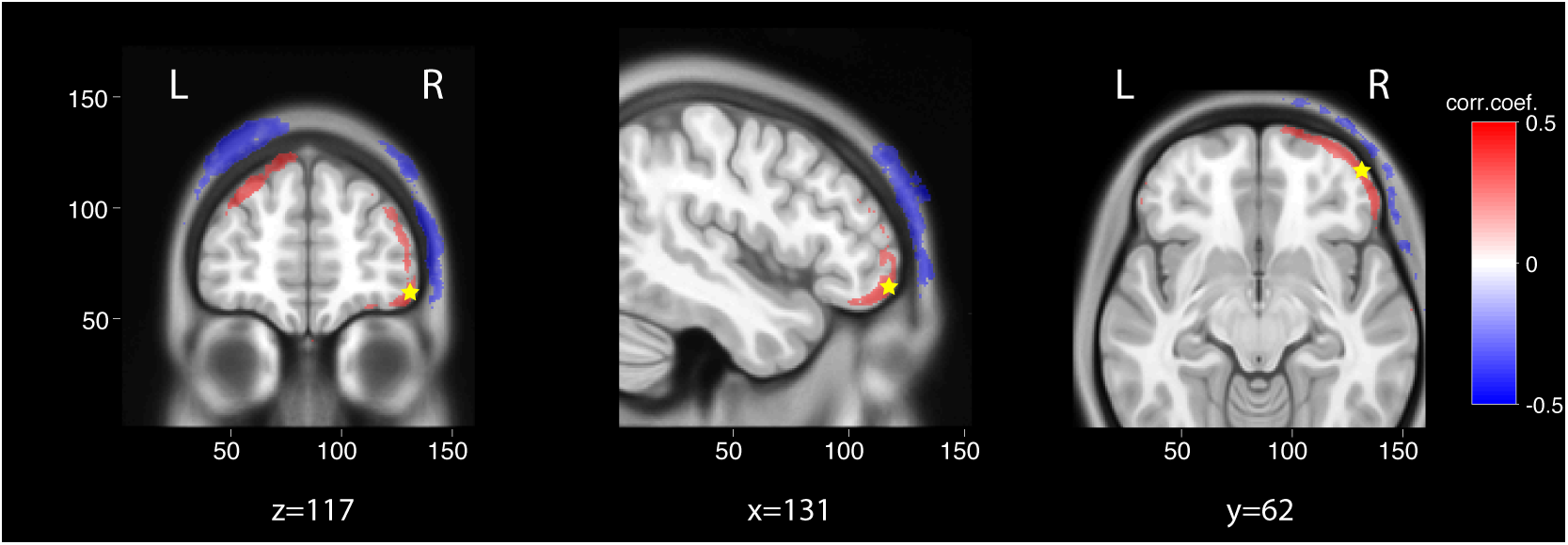
Correlation of electrophysiological response with electric fields magnitude. To determine the mechanism of action of tDCS, the pre/post change in EEG multivariate prediction of consciousness was correlated with tDCS-induced electric field distribution modeled on single-subject anatomy using T1-weighted MRI (n=47 patients). Correlations were restricted to areas with electric fields likely to have a physiological effect using a cut-off of mean electric fields>0.5 V/m. Multivariate EEG prediction of consciousness significantly correlated with higher electric fields in superficial cortical areas close to the stimulating electrodes (left dorsolateral prefrontal cortex and right supraorbital cortex) and with lower electric fields in the adjacent skin (voxel-wise significant positive (red) and negative (blue) correlations with p<0.01 uncorrected). The average correlation coefficient in the areas with were 0.433 and −0.432 in the areas with positive and negative correlations respectively. A statistical analysis on the strength of these mean correlations using 10000 permutations yielded p=0.0069 and p=0.0359 respectively (similar results were obtained when analysis was limited to areas with mean electric fields>1V/m, not shown).

### Discussion

In this study, we characterized the behavioral and electrophysiological responses to a single session of left-dorsolateral prefrontal cortex tDCS in the largest cohort of DoC patients to date. We first showed that behavioral improvement of consciousness was associated with an increase of resting-state putative markers of conscious state, together with the emergence of a neural signature of conscious access to auditory novelty. Then, we linked these electrophysiological changes to tDCS by showing a positive correlation between electrophysiological response and electric fields in prefrontal cortices, while a negative correlation was observed at the scalp.

In spite of the open-label design, several arguments strongly support a clinically relevant, genuine impact of tDCS. First, we replicated the response rate of a previous randomized controlled trial using the same stimulation parameters (21). Second, this response rate exceeded the changes associated with spontaneous fluctuations observed in the literature (40), and the short delay between evaluations (<2 hours) limited potential confounding factors that could account for the observed effects independently of tDCS (e.g. medications, sepsis,…). Lastly and most importantly, both the association of the behavioral response with an enhancement of EEG signatures of consciousness and the correlation of prefrontal cortices electric fields with the electrophysiological response are suggestive of a direct causal effect of tDCS on brain function, while the negative correlation at the scalp argues against indirect effects mediated by skin sensations. The use of such objective and quantified measures of brain activity is mandatory in transcranial electric stimulation studies, not only because it allows to investigate the causal link between behavior and cognitive function and the mechanisms through which tDCS acts, but also because proper blinding with a standard sham stimulation procedure seems difficult or even impossible to achieve (41).

The parallels between behavioral improvements and enhancement of putative EEG signatures of consciousness adds to our understanding of the neurophysiology of consciousness. Indeed, these results are the first to support the direct, if not causal, relationship between improvement of consciousness and increases in theta-alpha frequency power and functional connectivity (except for case reports with 4 patients (42); and vagus nerve stimulation in one patient (17)). So far, evidence was merely correlational, stemming from observational studies with inter-subject comparisons of separate populations of VS/UWS and MCS patients. Importantly, the findings from the present interventional study with intra-subject design and short inter-assessment delay indicate that these markers are not only specific to consciousness, but also sensitive to subtle behavioral improvements. Our study also emphasizes the central role of posterior associative cortices and posterior long-range functional connectivity in the theta band as key elements of consciousness (11), which could be the substrate for distant neural networks to generate coherent and sustained patterns of large-scale activity as proposed by the Global Workspace theory of consciousness (43, 44). Similarly, the all-or-none difference between R+ and R-during the oddball auditory paradigm supports the postulate identifying the P300 to a specific signature of conscious access (27, 45–47). This is also consistent with previous findings of an increase P300 amplitude in MCS patients after repetitive tDCS (48).

Our study also enriches the understanding of the mechanisms by which tDCS can elicit improvements of consciousness. Previous studies using fMRI in healthy subjects showed that anodal-tDCS over the left-dorsolateral prefrontal cortex could modulate the functional connectivity between prefrontal cortices and thalamus (49). Yet, very few studies have investigated the electrophysiological effects of tDCS in DoC patients. Some studies focused on differentiating tDCS impact based on the patients’ state of consciousness (50, 51). Others used a qualitative analysis of the EEG background activity to characterize tDCS response (24). Our study is the first to link electrophysiological changes with behavioral response to tDCS, and the first to link this to electric fields magnitudes in the brain. tDCS effects on the brain are thought to rely on direct modulation of neuronal excitability (34) or of synaptic plasticity (35) driven by the applied electric fields. However, some have proposed also indirect effect through the activation of peripheral nerves and in particular skin sensations (37). The association between electric fields magnitude with an objective EEG measure, unrelated to behavioral assessment, is a strong argument for an effect of tDCS in our population. Furthermore, the significant positive correlations we found between the electrophysiological response and electric fields over frontal cortical areas and negative correlations at the skin point to a cortical origin of the tDCS-related gains and against peripheral effect of skin sensations. The left-lateralized anterior topography of the P300 component, encompassing the anodal site of tDCS, also supports a relation between stimulation sites and enhancement of specific cortical networks. Finally, some R+ patients only improved their vigilance sub-score of the CRS-R, suggesting a cortically-mediated activation of the ascending reticular activation system. Together with the electric field distribution, the identification of specific spectral power and connectivity signatures of consciousness could pave the way for the development of more efficient stimulation tDCS strategies such as individually tailored electrode montages based on patient neuroanatomy. Indeed, the substantial heterogeneity in behavioral response to electrical stimulation (52) may originate from inter-subject variability in brain anatomy underlying stimulating sites (53). A recent study of prefrontal tDCS delivered during a decision-making task demonstrated that prefrontal cortical morphology differences between healthy subjects accounted for more than one third of the variability in tDCS efficacy (54). This may be even more critical in DoC patients who suffer from various and severe brain lesions (55–57), and in whom response to tDCS depends on residual brain metabolism and grey matter integrity (58). Our innovative modeling approach could herald the individual adaptation of transcranial electrical stimulation parameters to improve its therapeutic power on disorders of consciousness.

Our study is encouraging as it provides additional evidence in favor of an effect of tDCS in DoC patient together with the first mechanistic account of the improvement of consciousness by modulating residual cortical activity and cortico-cortical connectivity in this population.

### Methods

In this prospective case-control study, our main goal was to evaluate the impact of tDCS on brain activity (EEG) according to the presence/absence of behavioral response. In order to do that, we did a combined behavioral and electrophysiological assessment immediately before and after a single tDCS session (Figure 1A, see below).

#### Population

All patients referred to the Neurology Intensive Care Unit of the Pitié-Salpêtrière university hospital (Paris, France) for an evaluation of consciousness were screened for participation in the study. This standardized evaluation included detailed and repeated behavioral assessments, structural and functional brain-imaging recordings (standard and quantitative EEG, cognitive evoked potentials, MRI). In the absence of contra-indications to tDCS (pace-maker, metal in the head, uncovered craniectomy, refractory epilepsy), patients’ also had a single open-label tDCS session (following the procedure previously used by Thibault et al. (21) and described below). All non-comatose patients DoC patient, without mechanical ventilation nor contra-indication to tDCS were included. Consent was obtained from the patients’ relatives. The protocol conformed to the Declaration of Helsinki, to the French regulations, and was approved by the local ethic committee (*Comité de Protection des Personnes; CPP n° 2013-A01385-40*) Ile de France 1 (Paris, France) under the code *‘Recherche en soins courants’*.

#### Transcranial direct current stimulation (tDCS)

The stimulation consisted of a single open-label 20 minutes session of anodal tDCS stimulation over the left dorsolateral prefrontal cortex of 2 mA. The anode was placed over the left dorsolateral prefrontal cortex and the return electrode was placed over the right supraorbital frontal cortex (respectively F3 and Fp2 in the 10-20 international system EEG placement). Stimulation was delivered using Neuroelectrics Starstim 8 system ®, through 25 cm^2^ circular sponge electrodes soaked with saline solution. Impedance were kept below 10 kΩ during the whole stimulation session. We choose this montage to match previous studies of tDCS in DoC patients.

#### Behavioral evaluation

State of consciousness was assessed using the clinical gold-standard Coma Recovery Scale-Revised scale (CRS-R) which evaluates the presence or absence of responses on a set of hierarchically ordered items testing auditory, visual, motor, oromotor, communication and arousal function. CRS-R is both quantitative (scores range from 0 to 23) and qualitative with some key behaviors defining different states of consciousness (coma, VS/UWS, MCS or exit-MCS). Response to tDCS (R+) was defined a priori by an increase in CRS-R score after stimulation compared to the CRS-R score before stimulation by contrast with no change or a decrease in CRS-R score (R-). All CRS-R were performed by trained physicians (FF, BH and PP) at the same time of the day (end of the morning). Each patient was evaluated by the same physician, and physicians were not blinded to the intervention. However, it’s important to note that we used analyses based on objective EEG measures with the post-stimulation recording preceding the post-stimulation behavioral assessment, and that since all patients received the same active stimulation, the findings of the R+ vs. R-comparisons are unlikely to be explained by an expectation bias.

#### Electroencephalography analyses

##### Acquisition

The electrophysiological effect of tDCS were assessed using a 5 minutes resting-state EEG and an auditory oddball ERP paradigm, derived from the previously published local-global paradigm (27) designed to elicit automatic (mismatch negativity (59, 60) and P3a) and conscious (P3b) signatures of the detection of an auditory novelty (28, 29) (*SI Appendix A1*). Scalp EEG were recorded at a sampling rate of 250 Hz using a NetAmps 300 Amplifier (Electrical Geodesics, Eugene, Oregon) with a high-density sponge based 256 channels HydroCel Geodesic Sensor Net (Electrical Geodesics) referenced to the vertex. Importantly, the EEG cap was left in place throughout the tDCS session (stimulation electrodes were slithered underneath the EEG net) and before each recording, impedances were set below 100 kΩ.

##### Preprocessing

EEG data were processed using an automatized and hierarchical pipeline for artefact removal and extraction of EEG-measures previously described (13, 15, 26). Written in Python, C, and bash shell scripts and based on open source technologies, including the software MNE (61), the preprocessing workflow proceeded as follows:

EEG recordings were band-pass filtered (using a Butterworth 6th order high pass filter at 0.5 Hz and a Butterworth 8th order low pass filter at 45 Hz) with 50 Hz and 100 Hz notch filters. EEG during task were cut into epochs according to the onset of the fifth sound (800 ms before and 740 ms after) and resting-state EEG into 800 ms epochs with a 550 to 850 ms random jitter in-between (since this step is random, the preprocessing was repeated 100 times and subsequent measures were averaged over the 100 iterations). Channels that exceeded a 150 μV peak-to-peak amplitude in more than 50% of the epochs were rejected. Channels that exceeded a z-score of 4 across all the channels mean variance were rejected. This step was repeated two times. Epochs that exceeded a 150 μv peak-to-peak amplitude in more than 10% of the channels were rejected. Channels that exceeded a z-score of 4 across all the channels mean variance (filtered with a high pass of 25 Hz) were rejected. This step was repeated two times. The remaining epochs were digitally transformed to an average reference. Rejected channels were interpolated. EEG were deemed to pass this preprocessing step if at least 75% of the channels and at least 30% of the epochs were kept. To allow for the assessment of the electrophysiological effects of tDCS, only sessions in which both before and after stimulation recordings (either resting-state EEG or EEG during the task) passed the preprocessing stage were included in the analysis. For all subsequent EEG analyses, we only kept scalp electrodes (224 channels over 256).

##### Resting-state EEG analyses

###### Quantitative markers

Seventeen quantitative markers in three different domains were derived from the resting-state EEG recordings as in Sitt et al. (13) and Engemann et al. (15):

- Spectral domain:

Power spectrum density in each frequency band (δ: 1 – 4 Hz; θ: 4 – 8 Hz; α: 8 – 12 Hz; β: 12 – 30 Hz; γ: 30 – 45 Hz) were computed using Fast Fourier Transformation with the Welch method with a periodogram of 512 ms with 400 ms overlap. Raw and normalized spectral power (the sum of power in a frequency band reported to the power on all frequency bands of the spectrum sum) are reported for each frequency band.

Spectral entropy (characterizing the complexity of the spectrum), median spectral frequency, spectral edge 90th and 95th were computed.

- Connectivity:

Functional connectivity was assessed using the weighted symbolic mutual information (wSMI). This metric, able to capture non-linear coupling between pairs of electrodes, was introduced by King at al. (11) and reflects the statistical dependence of the transformation of the EEG signal into patterns of k discrete symbols (here k=3) sampled a different time interval (τ) which determines the frequency range specificity. In this study, we focused on the wSMI in the theta-alpha range (4-10 Hz, τ=32 ms), that we called wSMI θ, which has the best discriminative power across different state of consciousness (from VS/UWS to conscious subjects).

- Complexity:

Kolmogorov-Chaitin complexity and permutation entropy in the theta band, which computes the entropy of a signal transformed into discrete symbols.

All the markers were first computed at the single subject level: a value was obtained for each epoch at each channel (or channels pairs for the wSMI). Values were then averaged over the epochs using the trimmed mean 80% (mean of the distribution after trimming the 10% lowest and 10% highest values, a robust estimator of central tendency (62) to obtain a two-dimensional topographical representation over the 224 scalp electrodes for each subject. As the wSMI quantifies the shared information between electrodes, it was computed at each pairs of scalp electrodes (224X(224-1)/2 = 24976) and represented in a three-dimensional space. In addition, a two-dimensional representation was also obtained by resuming the value at each electrode by the median value of wSMI between one electrode and all the others. This averaging is closely related to the degree measure of the network in graph theory and highlights the sensors that have the strongest connections with other sensors, thus identifying hubs of connections. All analyses were done at the group level, using the mean across groups.

###### Multivariate Pattern Analysis (MVPA)

In addition to the effect on single markers, we assessed the overall effect of tDCS on brain activity using a MVPA approach as previously described by Sitt et al. (13). We used a linear Support Vector Classifier to predict the MCS-diagnosis (as opposed to VS/UWS) from the resting-state EEG derived markers. To that end, the 17 previously described markers were summarized using all four combination between their averages and fluctuations over time (trimmed mean 80% and standard deviation over the epochs) and space (mean and standard deviation over the scalp) resulting in 68 features for each subject. The support vector classifier algorithm using these features was trained against a previously published database of 142 EEG recordings (68 MCS and 75 VS/UWS) in 98 different patients (13, 15) with a 20% features selections (best 20% features on univariate analysis using F-tests) and 5-fold stratified cross-validation with a penalization parameter C, chosen by nested cross-validation among the values = [10^-6^ 10^-5^ 10^-4^ 10^-5^ 10^-4^ 10^-3^ 10^-2^ 10^-1^] using a grid-search method. The algorithm was then tested on each participants’ resting-state EEG to compute the classifier’s prediction of being MCS according to the brain activity both before and after the tDCS session separately. We used the Platt scaling method to obtain probabilities (between 0 and 1) from the classifier’s output. The change induced by tDCS was obtained by subtracting the probability when tested on the resting-state EEG before stimulation from the probability when tested on the resting-state EEG after stimulation. We used the scikit-learn software for machine learning (63).

##### Auditory oddball analyses

For the analysis of EEG during the auditory oddball paradigm, data were further low-pass filtered at 20 Hz and baseline corrected over the first 800 ms (from the beginning of the trial to the onset of the fifth sound).

###### ERP topographies

We first analyzed group-level event-related potentials (ERP) elicited by the auditory oddball paradigm. These were obtained by averaging trials of each condition with the trimmed mean 80% (across trial) over the scalp channels at the subject-level. We then computed group-level ERP to deviant minus standard trials, before and after tDCS, and compared R+ with R-(or VS/UWS with MCS & exit-MCS) with the following contrast: [[Deviant – Standard]after – [Deviant – Standard]before]R+ vs. [[Deviant – Standard]after – [Deviant – Standard]before]R-. Results are reported by the mean +- standard error of the mean.

###### Temporal generalization decoding

As classical ERP analyses are prone to a high inter-subject spatial variability, especially in brain-lesioned patients, we completed the topographical analyses with the analysis of the temporal dynamic of the ERP. To that end, with used the Temporal Generalization decoding method described by King et al. (30). This MVPA approach relies on the training of a classifier at each time point of the trial to distinguish deviant from standard trials at the single-subject level. Each classifier was then tested not only on the time sample it was trained on, but also on every other samples of the trial to see if decoding performances generalized in time. This procedure thus allows to extract patterns of brain activations associated with different cognitive tasks based on their temporal dynamics. This procedure has previously been applied to an auditory oddball paradigm and showed that the violation of auditory regularity led to two kinds neural activation patterns: an automatic, early and short-lived response to auditory novelty and a late and sustained (200-700 ms) pattern associated with the conscious access to the auditory novelty. This latter pattern probably reflects the same processus as the one engaged in the generation of the P300 during the same paradigm (33). Here we applied this MVPA procedure using linear support vector classifier (with 10 iterations of a stratified 5-fold cross-validation). Performances of the classifiers were expressed as the area under the curve (AUC).

#### Correlation of electric fields with EEG analysis

In order to estimate field distributions in individual patients we follow standard practice, namely, first we segment the head volume into different tissue compartments, then place virtual electrodes, and then use finite-element modeling to estimate current flow in the entire volume. We then correlate these field magnitudes with the EEG measure and use permutation statistic to determine statistical significance on the strength of these correlations. These steps are described in detail in the following sections.

##### MRI acquisition

T1-weighted MRI images were acquired during the patient hospitalization in 56 out of the 60 patients of the study, on a 3T General Electric Signa system (Milwaukee, WI) with a varying number of slices (from 152 to 234) and voxel sizes (from 0.42188 to 0.4883 mm^3^). This variability was not an issue because MRIs were down-sampled to a common 1 mm^3^ isotropic resolution in the process of co-registering the individual brains with a common standard (MNI152 standard head, see below).

##### Tissue Segmentation

Segmentation of the head volume is based on the T1-weighted MRI of each patient. Segmentation distinguishes the following six tissue types: scalp, skull, cerebrospinal fluid (CSF), gray matter, white matter, and air cavities (e.g. sinuses). Due to the substantial abnormality of these patients’ anatomy, popular segmentation algorithms developed for normal heads (e.g., Unified Segmentation as implemented in SPM8, (64) often fail to accurately classify different tissues, especially the enlarged ventricles in the brain of these patients. Therefore, we adopted a machine learning approach. Specifically, a volumetric deep convolutional neural network (DCNN) was trained with hand segmentations of MRI’s from 37 ischemic stroke heads available from an unrelated project (65). The trained DCNN was used to segment the 56 MRIs. Since the DCNN requires the input data to have the same size, the MRIs were first registered and resampled to a common space with 1 mm^3^ isotropic resolution before entering the DCNN. This was done by using the co-register function in SPM8 (66). Post-processing of the segmentations was done with a dense 3D conditional random field (CRF) (67). This step makes final corrections to the output of the neural network by incorporating known morphological constraints. These constraints are hand-coded in the weights of the CRF, which penalize neighboring tissue categories that are unrealistic, for example, brain cannot be next to air. This ensures among other things, that brain tissue is surrounded by the CSF (68). For validation purposes, two MRIs of the present patient population with prominent enlarged ventricles were manually segmented. We find that improved performance in terms of Dice score for the DCNN as compared to SPM8 (*SI section B6*, Figure S6). In the end, 47 MRI could be successfully segmented using this procedure and 9 had to be discarded due to insufficient quality (mainly because of movement artifacts).

##### Current flow modeling

The segmented tissues output from the DCNN were feed into an open-source software ROAST (69) to model the current-flow in the entire head volume. Electrodes were placed automatically by ROAST at positions F3 and Fp2 on the scalp. Electrodes were modeled as a disc with 25 cm^2 cross-sectional area and 2 mm thickness. Finite element meshing and solving were subsequently performed in ROAST fully automatically. Boundary conditions were set as 2 mA current injected at electrode F3, and 2 mA flowing out of electrode Fp2. Literature tissue conductivity values were used (70). To facilitate the voxel-level correlation analysis that is described in the next section, we registered and warped the electric field magnitude in each subject to the MNI152 standard head (version 2009a, (71, 72)). This was done by using the Unified Segmentation function (64) together with the DARTEL toolbox in SPM8 (73). Specifically, Unified Segmentation was run on the MRI of each subject, using the MNI152 standard head as the reference, during which a warping field from individual MRI to the standard head was estimated. The DARTEL function was then used to apply the estimated warping field on the electric field magnitude output from ROAST. In this way the field magnitude in all the subjects are co-registered with the voxel space of the standard head. The advantage of DARTEL over conventional affine transform is that it is capable of applying different warping field locally to different part of the image volume, and thus transforming different head shapes in each subject to a common shape represented by the standard head. This detailed alignment is important as fields differ drastically between cortical surface, CSF, bone and skin (due to very different conductances). Since these structures are all very thin, incorrect alignment results in a mismatch of tissue types across subjects, with potentially larger errors when correlating the EEG regressor variable with field magnitudes across subjects.

#### Statistical analyses

We first assessed the effect of prefrontal tDCS on both the behavior and a set of comprehensive and quantified electrophysiological markers of consciousness. To that end, we compared the EEG-derived markers and on the ERP according to the behavioral response to tDCS. To further understand the origin of significant differences if any, we also tested the after vs. before difference in each group separately when appropriate. The same types of analyses were performed contrasting tDCS-induced effect according to the state of consciousness before stimulation (VS/UWS vs. MCS and exit-MCS). Lastly, to understand the origin of these electrophysiological changes after tDCS, we correlated electric fields modeled on individual patients’ brain anatomy with the EEG multivariate prediction of consciousness.

##### Behavioral analysis and population characteristics

Population baseline characteristics (age, sex, time since brain injury, etiology and CRS-R scores) between R- and R+ and between VS/UWS and MCS & exit-MCS were compared using a Mann-Whitney-U for continuous data and the χ2 test for categorical variables as appropriate. In order to assess if there were differences in EEG data preprocessing between responders and non-responders, we used a non-parametric factorial analysis of repeated measure design. This procedure consists in a standard repeated measure analysis of variance (ANOVA) on aligned and rank transformed data (74) allowing for a more accurate assessment of the interaction term than with standard rank transformation. In this mixed design 2 by 2 design, we tested the effects of the behavioral response (between-subject factor: responder or non-responder) and tDCS stimulation (within-subject factor: before and after tDCS) on the number of channels/epochs rejected (dependent variable).

##### Electrophysiological data

For the analysis of multivariate MCS prediction, we used the same non-parametric factorial analysis with the behavioral response as between-subject factor and tDCS stimulation as the within-subject factor (before and after tDCS). All others resting-state EEG and auditory paradigm EEG analyses involved comparisons at multiple sensors and/or timepoints requiring a control procedure through permutations. However, there are debates on how permutations should be done in such a factorial design. In order to obtain a simpler contrast for the interaction term, we resumed the intra-subject effect of stimulation to the after minus before subtraction: [post – pre] in R+ vs. [post – pre] in R-. This analysis allowed us to assess the interaction while robustly controlling for multiple comparisons through a cluster-based permutation approach. (75). This two-step robust non-parametric procedure is able to control for the multiple comparisons over sensors and time by acknowledging the dependence between spatially close electrodes and timepoints. Firstly, a test statistic between the two groups was computed at each sample, i.e. each channel for the EEG markers, each channel-timepoint pair for the ERP and each channels pair for the temporal generalization decoding. Respectively, spatial, spatiotemporal and spatio-spatial clusters were constructed using a connectivity matrix with the inclusion of the test statistic corresponding to a first step statistic p-value of 0.05. For each cluster identified, the sum of test statistic over sensors included in the cluster (the cluster mass) was computed. The second step relied on the creation of surrogate datasets by 10000 random permutations of the groups’ labels to construct the distribution of the cluster mass under the null hypothesis that data from both groups are drawn from the same distribution. This allowed to estimate the probability of observing randomly each cluster constructed with the genuine data. As the first step statistic, we used a Welch test (unequal variance t-test, t-statitic) for the resting-state EEG markers analyses and the ERP topographies to compare mean voltage and a Mann-Whitney-U test (z-statistic) for multivariate analyses to compare median AUCs. For the additional analyses testing differences between after and before stimulation separately in each group, we used a dependent t-test and Wilcoxon Signed-Ranked test respectively. A type I error of 5% considered as significant. When present, we reported clusters’ effect size, defined as the effect size computed after averaging over the electrodes and/or time-points constituting the cluster in both populations. For the resting-state EEG an ERP topographies, we used the Hedges’ g coefficient (76), which is an approximation of Cohen’s d coefficient less prone to upward bias for small sample size. For the multivariate analyses, we reported the effect size measure *r:* 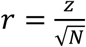. Where *z* is the z-statistic of the Mann-Whitney-U or Wilcoxon test and *N* the size of the population (77) *(SI Appendix A2)*. All effect sizes are reported with their bootstrapped 95% confidence interval computed using 10000 iterations.

##### Correlation of electric fields with EEG multivariate prediction of consciousness

To better characterize the potential mechanisms of action of tDCS in our cohort, we correlated electric fields magnitude distribution to the pre/post difference in multivariate prediction of consciousness. We chose this parameter for several reasons. First, the electrophysiological assessment was entirely automated and thus not susceptible to experimenter bias, in contrast to the behavioral assessment by the clinician who was not blinded. Second, the behavioral assessment with the CRS-R scale only gave a binary answer, with improvements in 9 “responders”. This small number of binary outcomes are not adequately powered to perform a whole-brain statistical analysis, in contrast to the graded outcomes of the electrophysiological assessment that is available for all 47 patients. Third, it is quite conceivable that there are neurophysiological changes that are not detected with the clinical behavioral assessment. Lastly, among all electrophysiological markers computed, we chose the MVPA prediction of consciousness because it systematically outperformed single EEG markers in predicting patients state of consciousness in previous studies (15). The individual subjects pre/post differences were correlated with the individuals’ electric fields in each voxel with mean electric field magnitude >0.5 V/m using Pearson correlation coefficients. This cut-off of electric field magnitude of at least 0.5 V/m in the average over 47 subjects was chosen based on fields magnitudes that are considered necessary to affect neural function (usually 1V/m or higher (39), though some effects were reported with fields as low as 0.25 V/m (38); at a mean of 0.5V/m some subjects will have weaker, other subjects stronger electric fields in those areas). For the uncorrected statistics we used a cut-off of p=0.01, corresponding to a correlation coefficient of 0.327 for positive correlations and −0.327 for negative correlations (N=47 patients). To quantify the overall strength of these correlations and the probability of observing it by chance we took the mean of the correlation coefficients in those areas and used shuffle statistics with 10,000 random permutations of patients’ labels. Note that the computational estimates of electric fields vary smoothly within a given tissue, such that correlation of one voxel naturally results in correlation of many neighboring voxels. Thus, extended areas of correlation (above the 1% threshold) are expected even for the null hypothesis of no correlation. As a result, size of the areas or the integral of the correlation over the area is not an appropriate metric. This motivated us to base the shuffle statistic more directly on the mean correlation coefficient as a measure of the strength of correlation.

##### Softwares

All statistical analyses were performed with python using *scipy (78), mne-python (61)* and *scikit-learn* (63) packages, except for the non-parametric factorial analysis performed in R statistical software with the *ARTool* (79) package and electric fields analysis performed in Matlab software. Throughout all EEG analyses, from preprocessing to statistical analyses, we tried to comply with the CODIBAS-MEEG Best Practices in Data Analysis and Sharing in Neuroimaging using MEEG initiative (80).

#### Data availability

The data that support the findings of this study are available from the corresponding author, upon reasonable request.

## Acknowledgements

This work was supported by : « Institut National de la Santé et de la Recherche Médicale » (BH, LN, JS), Sorbonne Université (LN), the James S. McDonnell Foundation (LN), FRM 2015 (LN), UNIM (LN), Académie des Sciences-Lamonica Prize 2016 (LN). The research leading to these results has received funding from the program “Investissements d’avenir” ANR-10-IAIHU-06 and by ANR grant “CogniComa” (ANR-14-CE-0013-03). BH was funded by « Poste d’Accueil Inserm » grant.

