## Supplementary Material for "Combined behavioral and electrophysiological evidence for a direct cortical effect of prefrontal tDCS on disorders of consciousness"

**A. Methodological considerations**

1. **Auditory oddball paradigm**
2. **Effect size measures**

**B. Supplementary results**

1. **Population description (Figure S1, Table S1 and Table S2)**
2. **Other resting state EEG markers according to tDCS response (Fig S2)**
3. **Resting state EEG markers according to the state of consciousness before stimulation (Fig S3)**
4. **Auditory oddball paradigm according to the state of consciousness before stimulation (Fig S4)**
5. **Auditory oddball paradigm in each groups (Fig S5)**
6. **Segmentation procedure example (Fig S6)**

**C. References**

### A. METHODOLOGICAL CONSIDERATIONS

#### 1. Auditory oddball paradigm

The auditory oddball paradigm was a modified short version of the previously published local-global paradigm (1) designed to elicit automatic (mismatch negativity (2, 3) and P3a) and conscious (P3b) signatures of the detection of an auditory novelty (4, 5).

Trials consisted of series of 5 sounds (50 ms high (A) or low pitch (B) complex tones, separated by 150 ms) with an inter-trial interval of 1350 to 1650 ms. The first four sounds were always identical. The fifth sound was the same in 80% of the trials (standard trials, AAAAA or BBBBB) and from a different pitch in 20% of the trial (deviant trials AAAAB or BBBBA). Standard and deviant trials were pseudorandomized with a total of 26 deviant trial. The experiment consisted of 2 blocks, with each block defined by the pitch of the standard tone (block 1: 80% AAAAA/20% AAAAB; block 2: 80% BBBBB/20% BBBBA). Before each block, patients were stimulated vocally and instructed to listen carefully to the sound and count the deviant trials.

#### 2. Effect size measures

For the resting-state EEG an event-related potentials topographies topographies, we used the Hedges's g coefficient (6), which is an approximation of Cohen's d (7) coefficient less prone to upward bias for small sample size :

$$g = d \times \left(1 - \frac{3}{4 \times (df) - 1}\right)$$

Where *df* is the number of degrees of freedom ($df = n1 + n2 - 2$) and *n1* and *n2* the respective number of patients in each population and d is Cohen's d:

$$d = \frac{\overline{x1} - \overline{x2}}{s^*}$$

Where $\overline{x1}$ and $\overline{x2}$ are the mean of both populations and $s^*$ the pooled standard deviation:

$$s^* = \sqrt{\frac{\sum((x1 - \overline{x1})^2 + (x2 - \overline{x2})^2)}{df}}$$

For the non-parametric analyses (CRS-R comparisons, comparisons of AUC and multivariate analyses), we report the effect size measure *r* (7):

$$r = \frac{z}{\sqrt{N}}$$

Where $z$ is the z-statistic of the Mann-Whitney-U or Wilcoxon test and $N$ the size of the population and the number of paired samples respectively.

### B. SUPPLEMENTARY RESULTS

#### 1. Population description

#### Flow chart

Among the 110 patients suffering from disorders of consciousness (DoC) that we assessed, 69 were eligible (33 were for the study and tDCS was delivered to 66 of these patients. Two patients were discarded before the preprocessing stage: one had a short and isolated epileptic seizure just after the tDCS session, and the other patient was in comatose state at the time of the recording (the Coma Recovery Scale-revised (CRS-R) (8, 9) was equal to 2 [0-0-1-1-0-0]). Four additional patients were discarded because of poor EEG data quality (see below). In case of repeated assessments of consciousness during this 3-year period for a given patient, we only kept the tDCS session corresponding to the best behavioral state of consciousness prior to tDCS in order to limit the possible impact of spontaneous fluctuations that may overestimate the genuine effect of tDCS. In the end, a total number of 60 patients were included in the analysis (Fig S1).

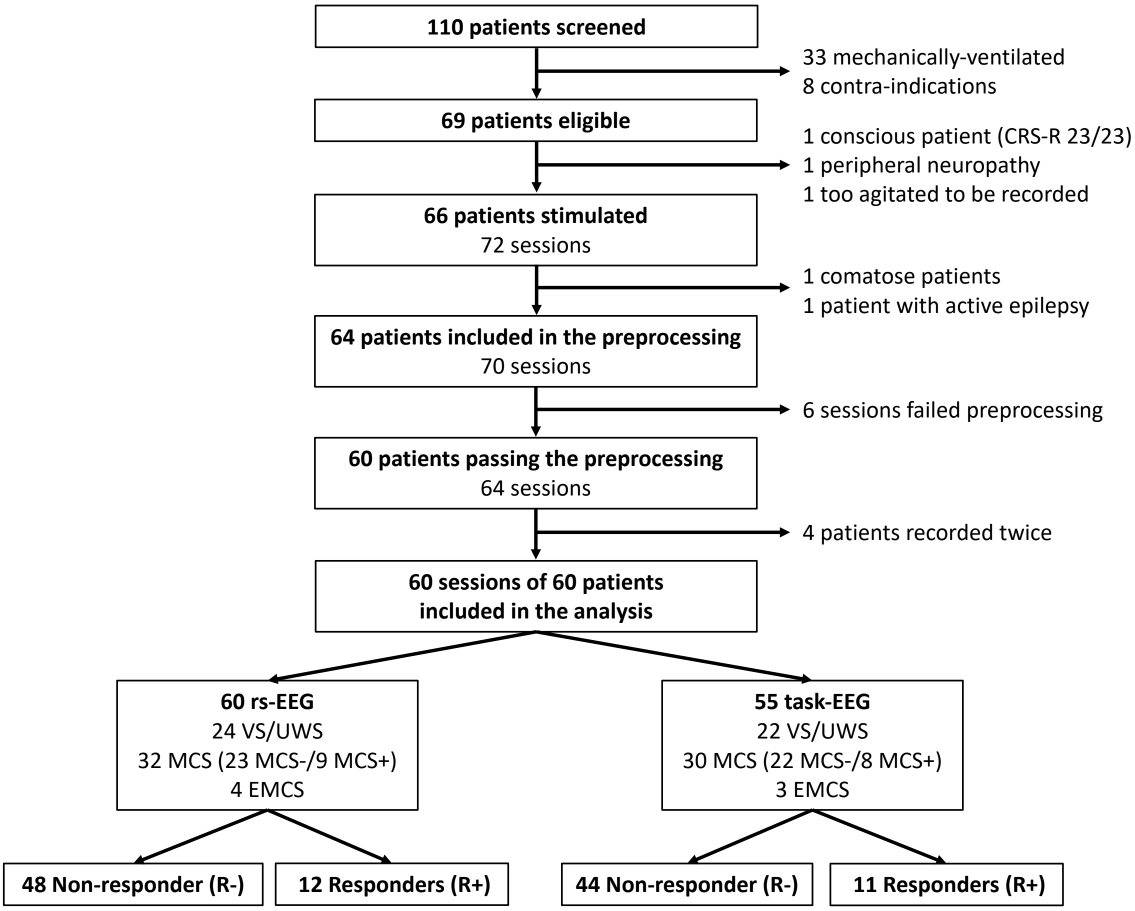

#### Figure S1. Flow chart

Flow chart showing the respective numbers of screened, eligible and included patients of the study together with the reasons for non-inclusion. The numbers of vegetative state/unresponsive wakefulness syndrome (VS/UWS), minimally conscious state (MCS), exit-minimally conscious state (EMCS), tDCS-responders (R+) and non-responders (R-) are presented for the two subpopulations in which resting state EEG (RS-EEG) and auditory oddball paradigm (task-EEG) were available

### Population characteristics

Individual patients' characteristics are presented in Table S1. Participants were typical DoC patients, with 24 VS/UWS, 32 MCS and 4 EMCS, with a median [interquartile range-IQR] age of 50 [32 ± 62] years, sex ratio of 1.3 and predominance of anoxo-ischemic (35%) and traumatic brain injury (29%). Median delay since brain injury was 195 [74-875] days. No significant differences were found between responders and non-responders in patients characteristics. Importantly, no differences were found neither in the pre-stimulation CRS-R arousal subscore, nor in the EEG preprocessing stage (Table S2).

#### Table S1. Patients characteristics

Patients demographic characteristics and behavioral evaluation (CRS-R total scores and subscores and state of consciousness) before and after stimulation, together with electrophysiological data available for the analysis (resting state and/or task-related EEG). Patients are sorted according to tDCS response (non-responders and responders), state of consciousness and then CRS-R score before stimulation.

**Abbreviations.** CRS-R: Coma Recovery Scale-revised; EEG: electroencephalogram; EMCS: exit minimally conscious state; F: female; M: male ; MCS: minimally conscious state; MRI: magnetic resonance imaging; Pat.: patients; R-: tDCS non-responder; R+: tDCS responder; rs: resting state; TBI : traumatic brain injury; tDCS: transcranial direct current stimulation; TSI: time since injury; VS/UWS: vegetative state/unresponsive wakefulness syndrome.

| Pat. # | Age | Sex | Etiology | TSI | CRS-R before | State before | CRS-R after | State after | EEG | MRI |
|---|---|---|---|---|---|---|---|---|---|---|
| **tDCS non-responders** | | | | | | | | | | |
| R-1 | 59 | F | Anoxia | 89 | 3 [0/0/1/1/0/1] | VS | 3 [0/0/1/1/0/1] | VS/UWS | rs, task | x |
| R-2 | 50 | M | Anoxia | 296 | 3 [0/0/1/1/0/1] | VS | 3 [0/0/1/1/0/1] | VS/UWS | rs | |
| R-3 | 70 | M | Anoxia | 31 | 3 [0/0/1/1/0/1] | VS | 3 [0/0/1/1/0/1] | VS/UWS | rs, task | x |
| R-4 | 63 | F | Stroke | 49 | 4 [1/0/1/1/0/1] | VS | 4 [1/0/1/1/0/1] | VS/UWS | rs, task | x |
| R-5 | 49 | F | Anoxia | 82 | 4 [1/0/1/1/0/1] | VS | 4 [1/0/1/1/0/1] | VS/UWS | rs, task | |
| R-6 | 31 | F | Anoxia | 2255 | 4 [1/0/1/1/0/1] | VS | 4 [1/0/1/1/0/1] | VS/UWS | rs | x |
| R-7 | 37 | F | Other | 76 | 4 [0/0/1/1/0/2] | VS | 4 [0/0/1/1/0/2] | VS/UWS | rs, task | x |
| R-8 | 61 | M | Anoxia | 60 | 5 [1/1/1/1/0/1] | VS | 5 [1/1/1/1/0/1] | VS/UWS | rs, task | x |
| R-9 | 70 | M | Stroke | 194 | 5 [0/1/2/1/0/1] | VS | 5 [0/1/2/1/0/1] | VS/UWS | rs, task | x |
| R-10 | 70 | M | Anoxia | 317 | 5 [2/0/1/1/0/1] | VS | 5 [2/0/1/1/0/1] | VS/UWS | rs, task | x |
| R-11 | 70 | M | Stroke | 114 | 5 [0/1/2/1/0/1] | VS | 5 [0/1/2/1/0/1] | VS/UWS | rs, task | x |
| R-12 | 22 | F | Anoxia | 39 | 5 [1/0/1/1/0/2] | VS | 5 [1/0/1/1/0/2] | VS/UWS | rs, task | x |
| R-13 | 68 | F | Other | 24767 | 5 [1/0/1/1/0/2] | VS | 5 [1/0/1/1/0/2] | VS/UWS | rs, task | x |
| R-14 | 53 | F | Anoxia | 2864 | 5 [0/1/2/1/0/1] | VS | 5 [0/1/2/1/0/1] | VS/UWS | rs, task | x |
| R-15 | 28 | M | Other | 100 | 5 [0/0/2/1/0/2] | VS | 5 [0/0/2/1/0/2] | VS/UWS | rs, task | x |
| R-16 | 24 | M | TBI | 620 | 6 [1/1/2/1/0/1] | VS | 6 [1/1/2/1/0/1] | VS/UWS | rs, task | |
| R-17 | 47 | M | Anoxia | 121 | 6 [1/1/2/1/0/1] | VS | 6 [1/1/2/1/0/1] | VS/UWS | rs, task | x |
| R-18 | 51 | F | Anoxia | 724 | 6 [1/0/2/1/0/2] | VS | 6 [1/0/2/1/0/2] | VS/UWS | rs, task | x |
| R-19 | 38 | M | TBI | 181 | 6 [1/1/1/1/0/2] | VS | 6 [1/1/1/1/0/2] | VS/UWS | rs, task | x |
| R-20 | 28 | M | TBI | 580 | 7 [2/0/2/1/0/2] | VS | 7 [2/0/2/1/0/2] | VS/UWS | rs, task | x |

| | | | | | | | | | | |
|---|---|---|---|---|---|---|---|---|---|---|
| R-21 | 32 | F | Anoxia | 172 | 5 [0/3/0/1/0/1] | MCS- | 5 [0/3/0/1/0/1] | MCS- | rs, task | x |
| R-22 | 51 | M | Other | 89 | 7 [0/2/3/1/0/1] | MCS- | 7 [0/2/3/1/0/1] | MCS- | rs, task | |
| R-23 | 27 | F | TBI | 374 | 7 [1/3/1/0/0/2] | MCS- | 7 [1/3/1/0/0/2] | MCS- | rs, task | x |
| R-24 | 42 | M | Stroke | 2069 | 8 [1/3/1/1/0/2] | MCS- | 8 [1/3/1/1/0/2] | MCS- | rs, task | x |
| R-25 | 67 | F | Other | 42 | 8 [1/3/1/1/0/2] | MCS- | 8 [1/3/1/1/0/2] | MCS- | rs, task | |
| R-26 | 34 | F | TBI | 681 | 8 [1/3/1/1/0/2] | MCS- | 8 [1/3/1/1/0/2] | MCS- | rs, task | x |
| R-27 | 42 | M | TBI | 1062 | 9 [1/3/2/1/0/2] | MCS- | 9 [1/3/2/1/0/2] | MCS- | rs, task | x |
| R-28 | 36 | M | TBI | 35 | 9 [1/0/5/2/0/1] | MCS- | 9 [1/0/5/2/0/1] | MCS- | rs, task | x |
| R-29 | 55 | M | Other | 1916 | 9 [1/3/2/1/0/2] | MCS- | 9 [1/3/2/1/0/2] | MCS- | rs, task | |
| R-30 | 30 | M | TBI | 110 | 9 [2/2/2/1/0/2] | MCS- | 9 [2/2/2/1/0/2] | MCS- | rs, task | x |
| R-31 | 35 | M | Anoxia | 500 | 9 [2/2/2/1/0/2] | MCS- | 9 [2/2/2/1/0/2] | MCS- | rs, task | x |
| R-32 | 55 | M | Other | 83 | 10 [0/3/5/1/0/1] | MCS- | 10 [0/3/5/1/0/1] | MCS- | rs, task | x |
| R-33 | 32 | M | TBI | 810 | 10 [2/0/4/2/0/2] | MCS- | 10 [2/0/4/2/0/2] | MCS- | rs, task | |
| R-34 | 63 | F | Anoxia | 58 | 10 [2/3/2/1/0/2] | MCS- | 10 [2/3/2/1/0/2] | MCS- | rs, task | x |
| R-35 | 22 | F | Other | 835 | 12 [1/3/5/1/0/2] | MCS- | 12 [1/3/5/1/0/2] | MCS- | rs, task | |
| R-36 | 36 | M | Other | 181 | 12 [2/3/3/2/0/2] | MCS- | 12 [2/3/3/2/0/2] | MCS- | rs | x |
| R-37 | 60 | F | Anoxia | 64 | 13 [2/3/4/2/0/2] | MCS- | 13 [2/3/4/2/0/2] | MCS- | rs, task | x |
| R-38 | 76 | M | Anoxia | 39 | 13 [2/3/3/3/0/2] | MCS- | 13 [2/3/3/3/0/2] | MCS- | rs, task | x |
| R-39 | 50 | M | Anoxia | 1678 | 14 [2/3/5/2/0/2] | MCS- | 14 [2/3/5/2/0/2] | MCS- | rs, task | |
| R-40 | 19 | F | Other | 142 | 14 [2/3/5/2/0/2] | MCS- | 14 [2/3/5/2/0/2] | MCS- | rs, task | x |
| R-41 | 18 | F | TBI | 1062 | 8 [3/1/2/1/0/1] | MCS+ | 8 [3/1/2/1/0/1] | MCS+ | rs | x |
| R-42 | 61 | M | Stroke | 50 | 8 [3/1/2/1/0/1] | MCS+ | 8 [3/1/2/1/0/1] | MCS+ | rs, task | |
| R-43 | 48 | M | TBI | 2917 | 11 [3/3/2/1/0/2] | MCS+ | 11 [3/3/2/1/0/2] | MCS+ | rs, task | x |
| R-44 | 25 | F | Stroke | 746 | 14 [3/3/5/0/1/2] | MCS+ | 14 [3/3/5/0/1/2] | MCS+ | rs, task | x |
| R-45 | 75 | F | TBI | 53 | 16 [2/3/5/3/1/2] | MCS+ | 16 [2/3/5/3/1/2] | MCS+ | rs, task | x |
| R-46 | 44 | F | Stroke | 1956 | 14 [2/3/6/1/0/2] | EMCS | 14 [2/3/6/1/0/2] | EMCS | rs, task | x |
| R-47 | 25 | M | TBI | 1105 | 21 [4/5/6/1/2/3] | EMCS | 21 [4/5/6/1/2/3] | EMCS | rs, task | x |
| R-48 | 71 | F | Other | 519 | 22 [4/5/5/3/2/3] | EMCS | 22 [4/5/5/3/2/3] | EMCS | rs, task | x |
| **tDCS responders** | | | | | | | | | | |
| R+1 | 67 | M | Anoxia | 196 | 4 [0/1/1/1/0/1] | VS | **5** [0/1/1/1/0/**2**] | VS/UWS | rs, task | x |
| R+2 | 57 | F | Other | 996 | 4 [1/0/1/1/0/1] | VS | **5** [1/0/1/1/0/**2**] | VS/UWS | rs, task | |
| R+3 | 68 | F | Anoxia | 52 | 5 [1/0/2/1/0/1] | VS | **6** [1/0/2/1/0/**2**] | VS/UWS | rs, task | x |
| R+4 | 22 | M | Anoxia | 354 | 5 [1/1/1/1/0/1] | VS | **8** [**2**/**2**/1/1/0/**2**] | **MCS-** | rs, task | x |
| R+5 | 74 | M | Other | 30 | 5 [0/0/3/1/0/1] | MCS- | **9** [0/**3**/3/1/0/1] | MCS- | rs, task | x |
| R+6 | 58 | M | Other | 67 | 13 [2/2/5/2/0/2] | MCS- | **14** [**3**/2/5/2/0/2] | **MCS+** | rs, task | x |
| R+7 | 30 | F | TBI | 643 | 13 [2/3/5/1/0/2] | MCS- | **16** [**3**/**5**/5/1/0/2] | **MCS+** | rs, task | |
| R+8 | 55 | M | TBI | 2089 | 11 [3/3/2/1/0/2] | MCS+ | **12** [3/3/2/1/**1**/2] | MCS+ | rs, task | x |
| R+9 | 65 | M | Anoxia | 180 | 12 [3/3/2/2/0/2] | MCS+ | **13** [**4**/3/2/2/0/2] | MCS+ | rs, task | x |
| R+10 | 32 | M | TBI | 2501 | 12 [3/3/3/1/0/2] | MCS+ | **19** [**4**/**5**/**6**/**1**/1/2] | **EMCS** | rs | x |
| R+11 | 54 | F | Stroke | 1341 | 20 [3/5/5/3/1/2] | MCS+ | **21** [3/5/5/3/**2**/2] | **EMCS** | rs, task | |
| R+12 | 29 | M | TBI | 59 | 19 [3/5/6/2/1/2] | EMCS | **20** [3/5/6/**3**/1/2] | EMCS | rs, task | x |

**Table S2. Resting state and Task-EEG baseline characteristics and preprocessing**

Population demographic, behavioral and preprocessing characteristics at baseline and EEG preprocessing characteristics. Statistical comparison between responders and non-responders were computed using Mann-Whitney-U test for continuous data, fisher exact test for qualitative data and non-parametric ANOVA for preprocessing data. No significant main effect of either tDCS or behavioral response nor interaction between the two was found for any of the preprocessing comparisons.

**Abbreviations.** CRS-R: Coma Recovery Scale-revised; EEG: electroencephalogram; EMCS: exit minimally conscious state; IQR: inter-quartile range; MCS: minimally conscious state; n: number; NS: not significant; R-: tDCS non-responder; R+: tDCS responder; rs: resting state; TBI: traumatic brain injury; tDCS: transcranial direct current stimulation; TSI: time since injury; VS/UWS: vegetative state/unresponsive wakefulness syndrome.

| RESTING STATE | | | | |
|---|---|---|---|---|
| **Variable** | **All**<br>*N=60* | **Responders**<br>*N=12* | **Non-responders**<br>*N=48* | **p-value** |
| **Age**, years, *median [IQR]* | 50 [32-62] | 56 [32-66] | 48 [32-61] | 0.44 |
| **Sex ratio** | 1.3 | 2 | 1.2 | 0.53 |
| **Time since injury**, days, *median [IQR]* | 195 [74-875] | 275 [65-1082] | 188 [81-816] | 0.85 |
| **Etiology**, *n (%)*<br>Anoxia<br>TBI<br>Stroke<br>Other | <br>21 (35)<br>17 (29)<br>8 (13)<br>14 (23) | <br>4 (33)<br>4 (33)<br>1 (9)<br>3 (25) | <br>17 (35)<br>13 (27)<br>7 (15)<br>11 (23) | 1.0 |
| **SoC before**, *n (%)*<br>UWS<br>MCS<br>EMCS | <br>24 (40)<br>32 (53)<br>4 (7) | <br>4 (33)<br>7 (58)<br>1 (9) | <br>20 (43)<br>25 (52)<br>3 (5) | 0.89 |
| **Pre-tDCS CRS-R**, *median [IQR]*<br>- Total score<br>- Arousal score | <br>8 [5-12]<br>2 [1-2] | <br>11.5 [5-13]<br>2 [1-2] | <br>8 [5-10]<br>2 [1-2] | <br>0.47<br>0.94 |
| **Preprocessging**, *median [IQR]*<br>- Nb of rejected channels<br>- before<br>- after<br>- Nb of rejected epochs<br>- before<br>- after | <br><br>16.9 [10.9- 21.7]<br>15.1 [10.5-23.2]<br><br>8.7 [2.9-21.5]<br>9.5 [2.3-27.1] | <br><br>14.7 [11.6-22.1]<br>14.9 [12.3-18.7]<br><br>8.7 [5.5-15.7]<br>9.5 [3.7-14.4] | <br><br>16.9 [10.7-20.4]<br>15.9 [10.3-25.4]<br><br>9.0 [2.4-22.7]<br>9.4 [2.2-29.6] | <br><br>NS<br><br><br>NS<br> |

| TASK-EEG | | | | |
|---|---|---|---|---|
| **Variable** | **All**<br>*N=55* | **Responders**<br>*N=11* | **Non-responders**<br>*N=44* | **p-value** |
| **Age**, years, *median [IQR]* | 51 [32-63] | 57 [42-66] | 48 [32-62] | 0.40 |
| **Sex ratio** | 1.3 | 1.8 | 1.2 | 0.74 |
| **Time since injury**, days, *median [IQR]* | 181 [66-778] | 196 [63-820] | 157 [67-740] | 0.95 |
| **Etiology**, *n (%)*<br>Anoxia<br>TBI<br>Stroke<br>Other | <br>19 (35)<br>15 (27)<br>8 (14)<br>13 (24) | <br>4 (37)<br>3 (27)<br>1 (9)<br>3 (27) | <br>15 (34)<br>12 (27)<br>7 (16)<br>10 (23) | <br>1.0 |
| **SoC before**, *n (%)*<br>UWS<br>MCS<br>EMCS | <br>23 (42)<br>28 (51)<br>4 (7) | <br>4 (36)<br>6 (54)<br>1 (11) | <br>19 (43)<br>22 (50)<br>3 (7) | <br>1.0 |
| **Pre-tDCS CRS-R**, *median [IQR]*<br>- Total score<br>- Arousal score | <br>8 [5-12]<br>2 [1-2] | <br>11 [5-13]<br>2 [1-2] | <br>8 [5-10]<br>2 [1-2] | <br>0.75<br>0.68 |
| **Preprocessging**, *median [IQR]*<br>- Nb of rejected channels<br>- before<br>- after<br>- Nb of rejected epochs<br>- before<br>- after | <br><br>19.0 [11.5- 27.0]<br>18.0 [12.5-29.0]<br><br>15.0[4.0-33.5]<br>9.0 [4.0-40.5] | <br><br>17.0 [12.0-20.5]<br>19.0 [14.0-25.5]<br><br>9.0 [1.5-14.5]<br>7.0 [2.0-12.0] | <br><br>19.5 [11.8-27.3]<br>17.0 [12.0-33.0]<br><br>18.5 [4.0-41.5]<br>14.0 [4.0-43.0] | <br><br>NS<br><br><br>NS<br> |

### 2. Other resting state EEG markers according to tDCS response (Fig S2)

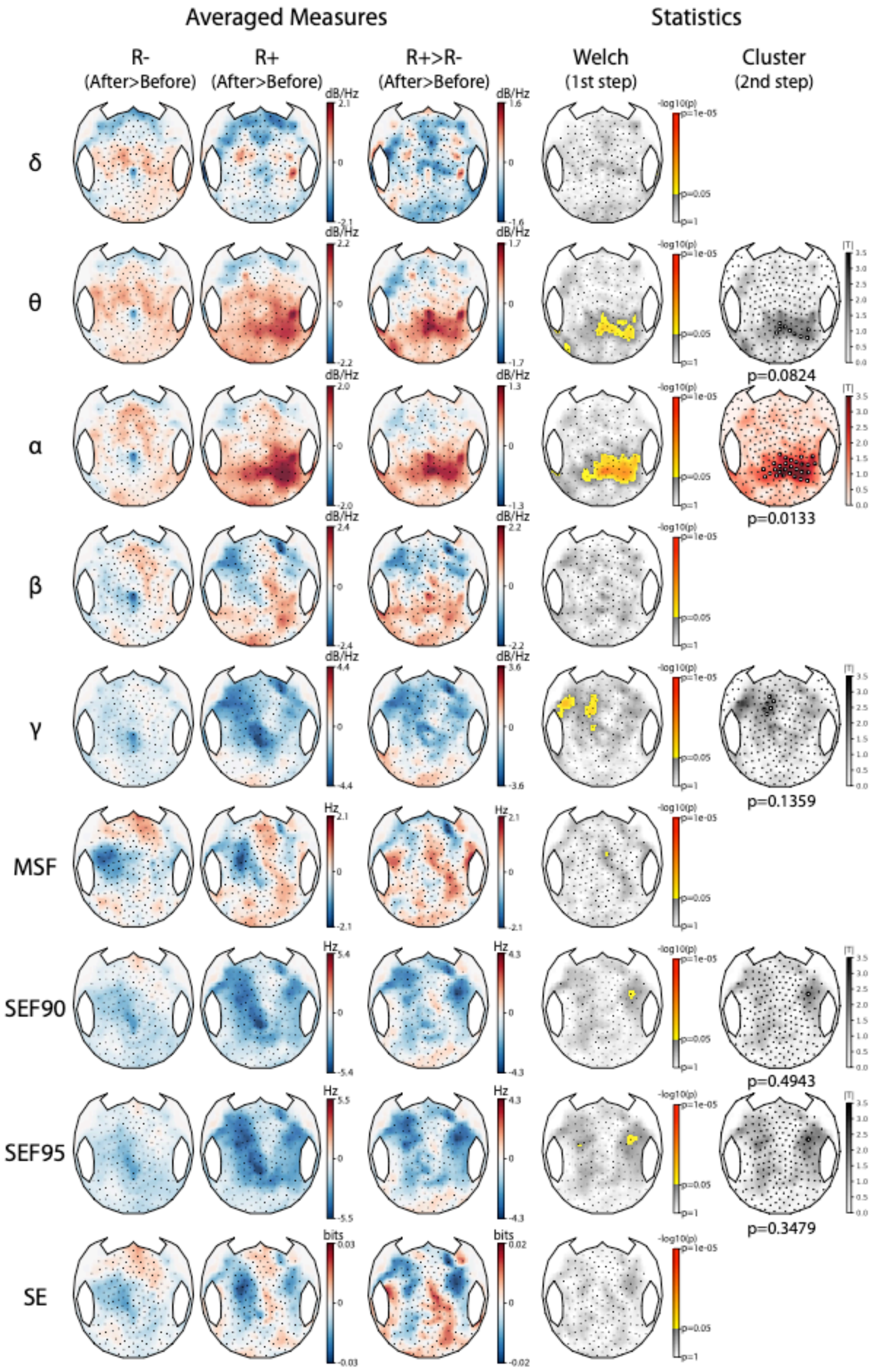


**Figure S2. Other resting state EEG markers according to tDCS response.** Topographical representations of the tDCS-induced changes in raw spectral power (delta δ, theta θ, alpha α, beta β and gamma γ), median spectral frequency (MSF), spectral edge frequency 90% and 95% (SEF90 and SEF95) and spectral entropy (SE) over the 224 scalp electrodes. After minus before differences are presented for both non-responders (R-) and responders (R+) (left columns), followed by the contrast between the two (middle columns) and the corresponding statistical comparison using a two-steps spatial cluster-based permutation approach (right columns). A significant centro-parietal cluster was found α power ($p = 0.0133$). Absolute t-values are plotted with a red color scale when a significant cluster was found and in grey otherwise. Electrodes forming the cluster are highlighted by white circles.

### 3. Resting state EEG markers according to the state of consciousness before stimulation (Fig S3)

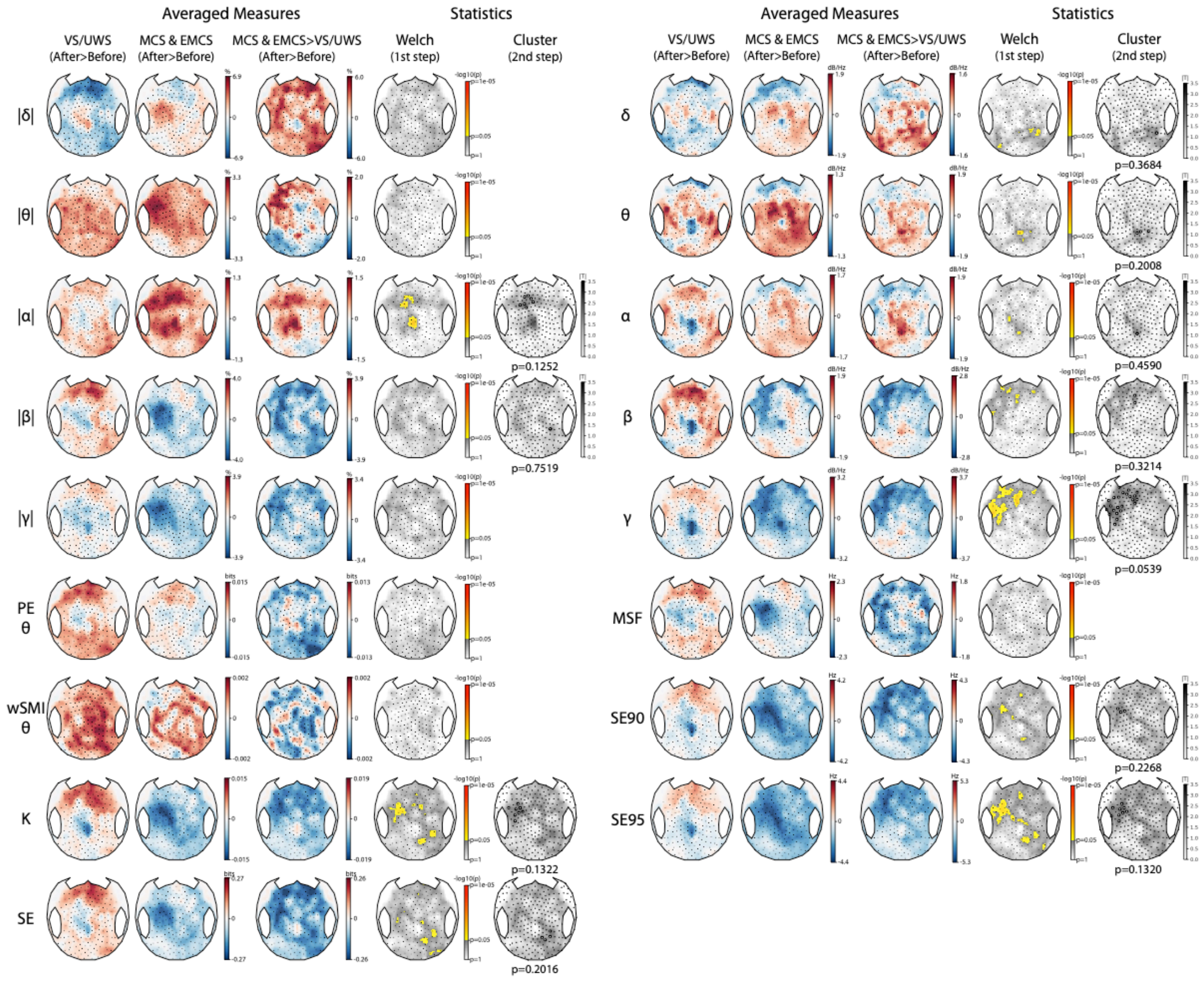


**Figure S3. Resting state EEG markers according to the state of consciousness before stimulation.** Topographical representations of the tDCS-induced changes in raw and normalized spectral power (delta δ, theta θ, alpha α, beta β and gamma γ), permutation entropy in the theta-alpha band (PE θ), weighted symbolic mutual information in the theta-alpha band (wSMI θ), Kolmogorov complexity (K), spectral entropy (SE), median spectral frequency (MSF) and spectral edge frequency 90% and 95% (SEF90 and SEF95) over the 224 scalp electrodes according to the state of consciousness before stimulation. After minus before differences are presented for both vegetative state/unresponsive wakefulness syndrome state (VS/UWS) and minimally and exit-minimally conscious state (MCS & EMCS) (left columns), followed by the contrast between the two (middle columns) and the corresponding statistical comparison using a two-steps spatial cluster-based permutation approach (right columns). No significant cluster was found for any of the studied EEG markers.

### 4. Auditory oddball paradigm according to the state of consciousness before stimulation (Fig S4)

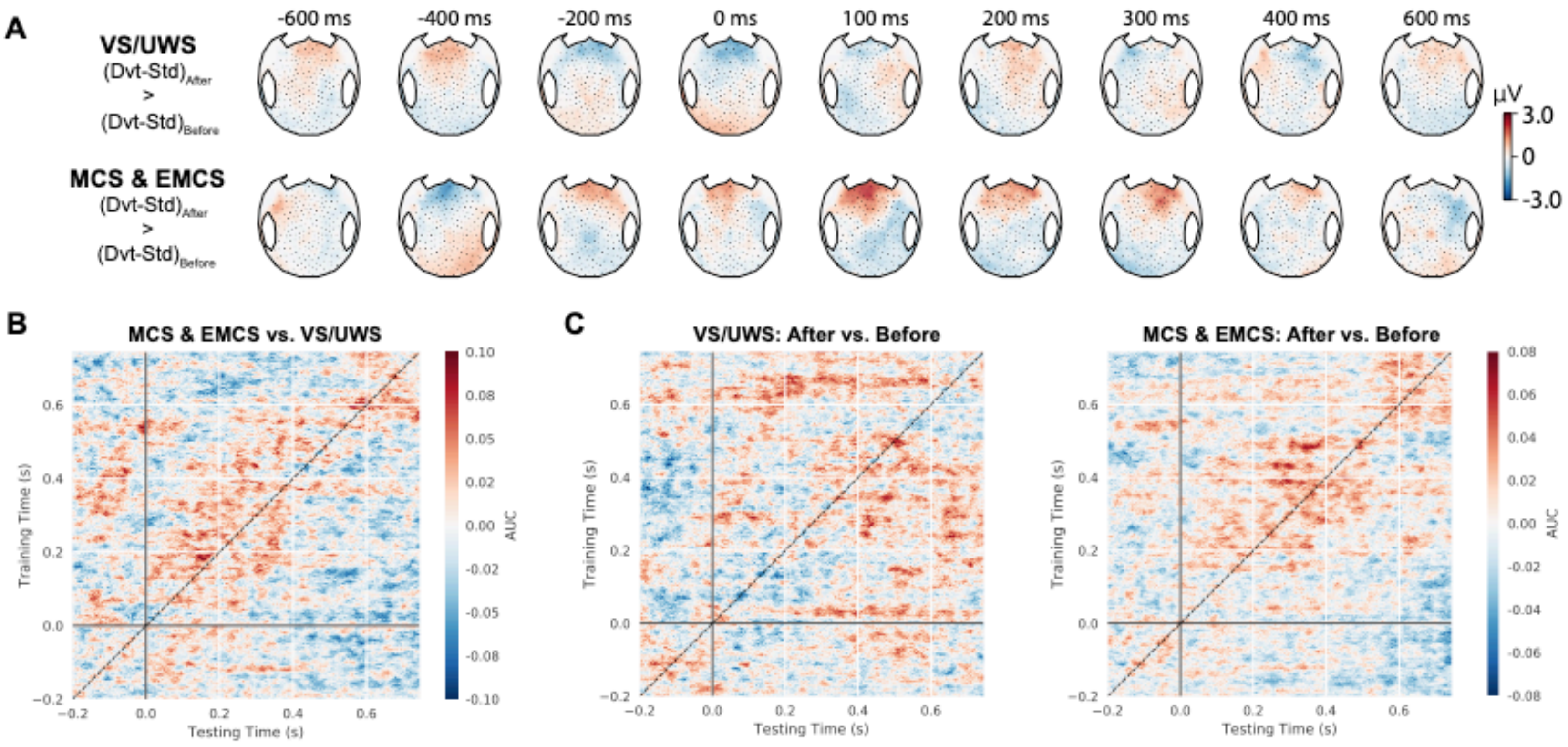


**Figure S4. Auditory oddball paradigm according to the state of consciousness before stimulation.**
(A). Topographical representation over time of the tDCS-induced changes in event-related potentials (After > Before difference of the Deviant (Dvt) > Standard (Std) contrast) during the auditory oddball paradigm in vegetative state/unresponsive wakefulness syndrome state (VS/UWS, top) and minimally and exit-minimally conscious state (MCS & EMCS, bottom). tDCS seemed to induce a more pronounced anterior positivity in the range of the P300 but no significant cluster was found when comparing both groups. (B) Temporal generalization decoding analysis showing the contrast in median AUC difference (after minus before) between VS/UWS and MCS & EMCS. Again, no significant difference between both groups was observed.

**5. Auditory oddball paradigm in each groups (Fig S5)**

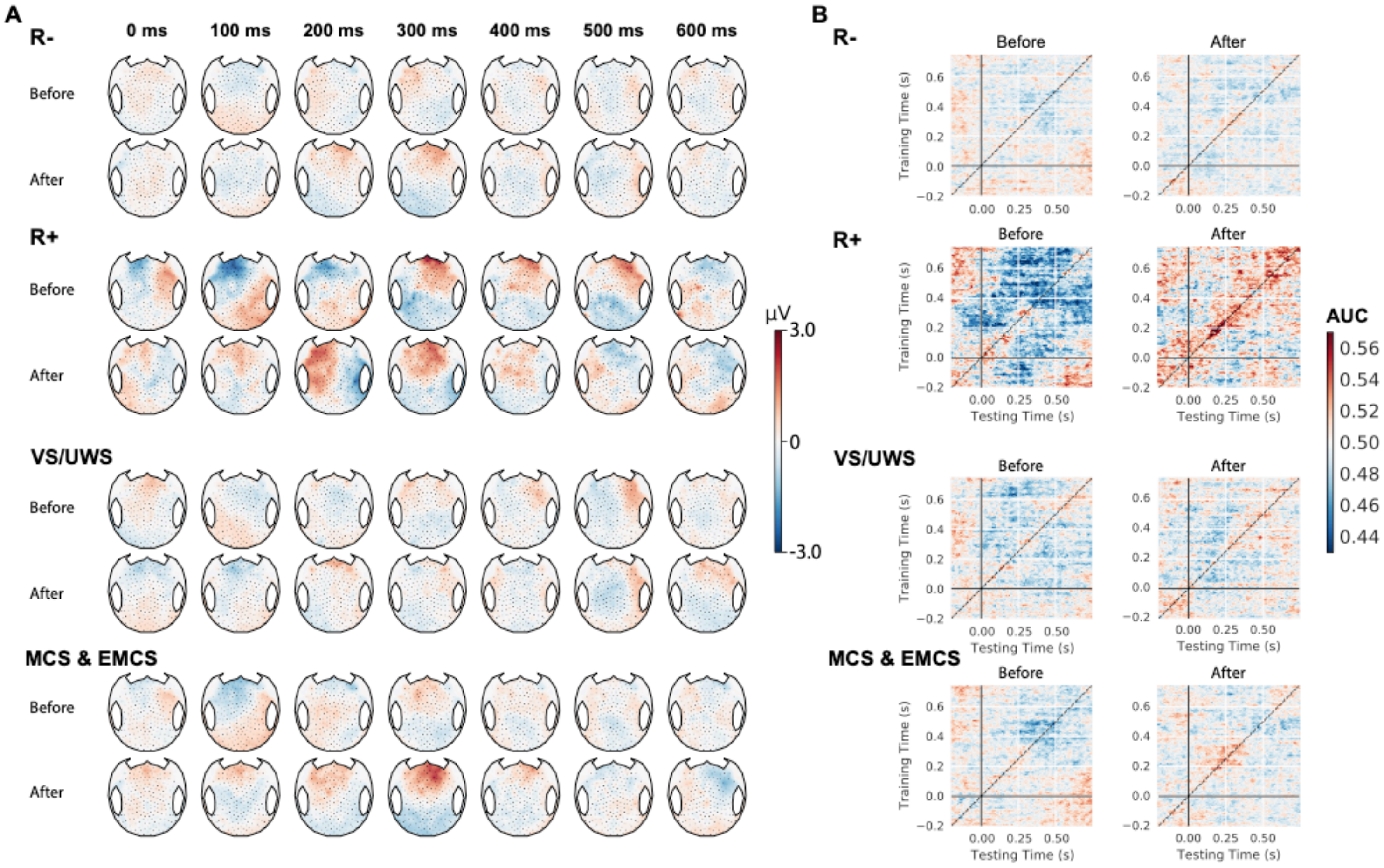


**Figure S5. Auditory oddball paradigm in each groups.**

(A). Topographical representation of the deviant (Dvt) > Standard (Std) event-related potentials during the auditory oddball paradigm before and after tDCS separately in non-responders (R-), responders (R+), vegetative state/unresponsive wakefulness syndrome state (VS/UWS) and minimally and exit-minimally conscious state (MCS & EMCS). (B) Corresponding temporal generalization decoding analysis for the Dvt vs. Std decoding.

### 6. Segmentation procedure example (Fig S6)

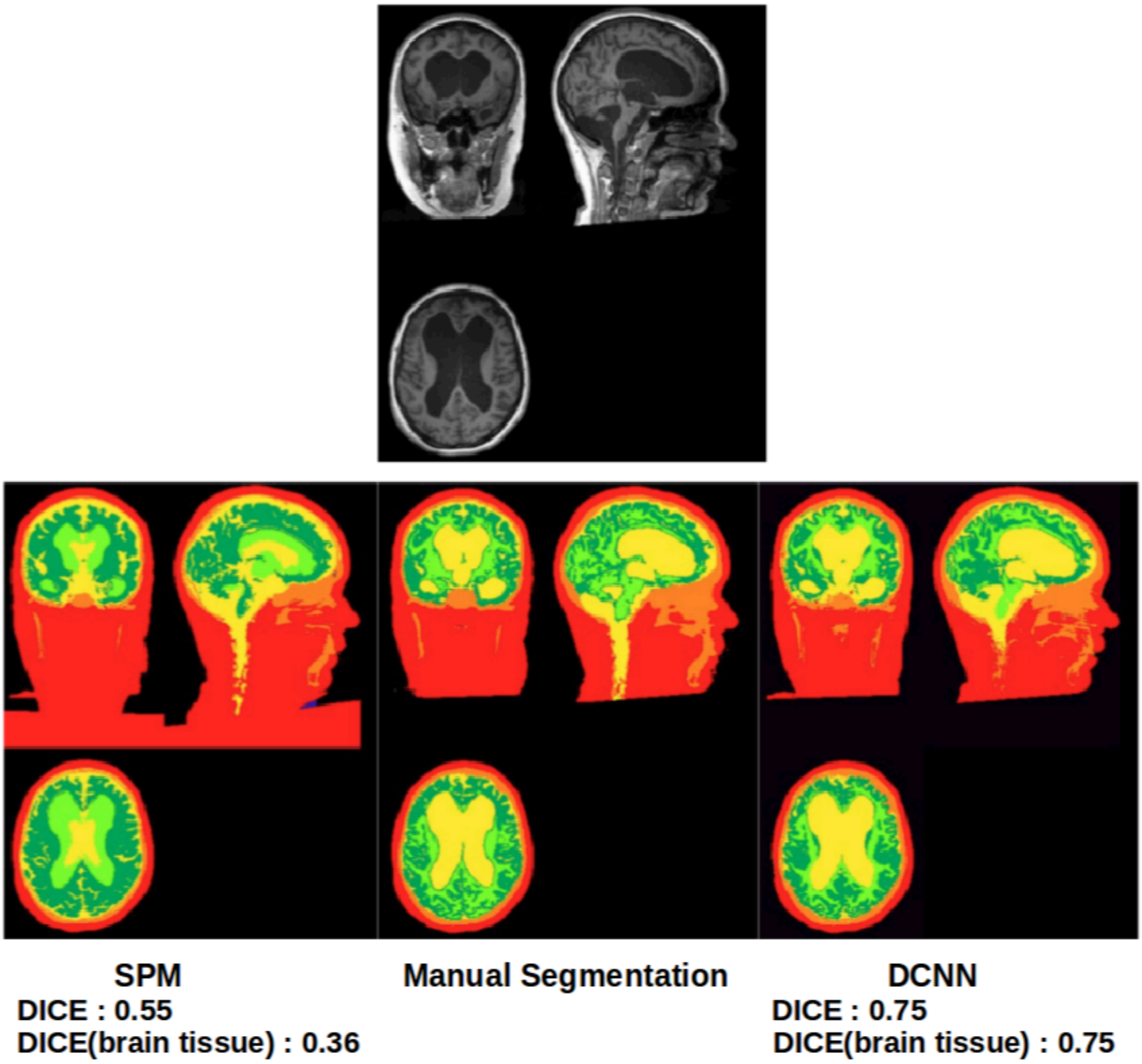


**Figure S6. Segmentation procedure example**

Segmented MRI using a deep convolutional neural network (DCNN) outperforms SPM on a subject with abnormal anatomy. When comparing the segmentations to manual segmentation, the neural network achieves a higher agreement as measured by the Dice score.
